# Spatiotemporally organized immunomodulatory response to SARS-CoV-2 virus in primary human broncho-alveolar epithelia

**DOI:** 10.1101/2023.03.30.534980

**Authors:** Diana Cadena Castaneda, Sonia Jangra, Marina Yurieva, Jan Martinek, Megan Callender, Matthew Coxe, Angela Choi, Juan García-Bernalt Diego, Jianan Lin, Te-Chia Wu, Florentina Marches, Damien Chaussabel, Peter Yu, Andrew Salner, Gabrielle Aucello, Jonathan Koff, Briana Hudson, Sarah E. Church, Kara Gorman, Esperanza Anguiano, Adolfo García-Sastre, Adam Williams, Michael Schotsaert, Karolina Palucka

## Abstract

The COVID-19 pandemic continues to be a health crisis with major unmet medical needs. The early responses from airway epithelial cells, the first target of the virus regulating the progression towards severe disease, are not fully understood. Primary human air-liquid interface cultures representing the broncho-alveolar epithelia were used to study the kinetics and dynamics of SARS-CoV-2 variants infection. The infection measured by nucleoprotein expression, was a late event appearing between day 4-6 post infection for Wuhan-like virus. Other variants demonstrated increasingly accelerated timelines of infection. All variants triggered similar transcriptional signatures, an “early” inflammatory/immune signature preceding a “late” type I/III IFN, but differences in the quality and kinetics were found, consistent with the timing of nucleoprotein expression. Response to virus was spatially organized: CSF3 expression in basal cells and CCL20 in apical cells. Thus, SARS-CoV-2 virus triggers specific responses modulated over time to engage different arms of immune response.

## Introduction

The COVID-19 pandemic is still ongoing despite development of preventive vaccines. SARS-CoV-2 virus first targets the epithelial cells of the respiratory tract ^1,2^ leading to a wide range of clinical presentations from asymptomatic to an uncontrolled inflammatory response with massive cytokine release and significant alveolar damage^1,3,4^. The airway epithelium constitutes a sophisticated barrier to maintain homeostasis and defend the lungs against pathogens^5^. Since the onset of the pandemic, several studies have looked at the pathways involved in the infection. To this end, angiotensin converting enzyme 2 (ACE2, reported as an interferon-stimulated gene (ISG))^6–8^ and transmembrane Serine protease-2 (TMPRSS2) have been shown to be major determinants for viral entry^9–11^. Recent studies have suggested additional coronavirus-associated receptors and factors such as Furin, CD147, CD209 and AXL^12^. Key gene signatures have been associated with the cytokine storm characterizing SARS-CoV-2 infection such as G-CSF, IL-6, CXCL8, CXCL10, IL1b, TNF, CCL20, CXCL1 and CXCL3^13,14^. Innate (neutrophils^3,15^, NK, macrophages^16–18^, dendritic cells^19,20^, complement^21–24^) and adaptative immune cells (T, B cells and antibodies)^25–27^, have been reported to play a critical role in SARS-CoV-2 infection, but their relative contributions and interplay with the lung epithelium are not yet fully understood. The rapid evolution of this virus and emergence of variants, such as newer Omicron subvariants, have displayed a potential increase in transmissibility, lower vaccine efficacy and an increased risk of reinfection necessitate a better understand of the early events that would determine the fate of the immune response.

Many studies have investigated SARS-CoV-2 variants to understand their transmissibility, disease, severity and/or ability towards immune escape in the hosts^28^. Several reports have stated the role of a single or multiple amino acid mutations in SARS-CoV-2 genome, especially in the Spike (S) region, in facilitating viral transmission and escape from neutralizing antibodies. One such example is N501Y mutation in the RBD of S protein, which enhances its binding to ACE2 receptor on host cells, resulting in an increased transmissibility and virulence^29–31^. Besides, the Delta variant displays an enhanced spike-mediated syncytia formation compared to the other SARS-CoV-2 variants, allowing an increased infectivity in lung epithelial cells^32,33^. For Omicron subvariants more than 30 mutations have been described in the Spike region resulting in altered tropism and reduced use of TMPRSS2 for entry/spread through cell-cell fusion^33,34^. Thus, Omicron (BA.1, BA.2) is capable of infecting mainly upper airway epithelium rather than alveolar epithelium, and form syncytia^33,35^. More recently emerged Omicron subvariants (BA.4; BA.5; BQ.1.1 and XBB.1.5) have been reported to exhibit an enhanced infectivity and antibody evasion capacity, even higher than BA.1 and BA.2 ^36–38^.

To better understand the early spatiotemporal events happening in lung epithelium during SARS-CoV-2 infection and their potential biological consequences, we based our study on an *in vitro* model of primary human airway epithelial cells: air-liquid interface (ALI) cultures generated from human lung organoids from healthy donors. Such cultures effectively recapitulate *in vivo* airway epithelium architecture^39,40^ and we validated their permissibility to viral infection by exposure to SARS-CoV-2 (Wuhan-like, USA/WA1-2020) and its variants (Beta, Delta and Omicron BA.1 subvariant). Our findings reveal spatiotemporal regulation of cytokines, chemokines and suggest that different cells in the lung epithelium may regulate different aspects of immune response to SARS-CoV-2. Furthermore, the use of lung organoid-derived ALI from a variety of donors revealed differential infection dynamics reflecting biological differences between individuals, potentially explaining some of the diverse clinical outcomes observed between patients.

## Results

### Establishing 3D differentiated primary human air-liquid interface cultures

We established epithelial lung organoids from viable cryopreserved lung fragments obtained from four donors (**Table S1**), using method described by Sachs et al.^41^ (**Figure S1A**). Lungs from two donors were acquired before the pandemic and the two others during the early stages of the pandemic (tested negative for SARS-CoV-2 by PCR). Organoids were stored viably frozen and used to generate ALI cultures^41^ (**Figure 1A**). Epithelial differentiation and barrier integrity was monitored by measuring the transepithelial electrical resistance (TEER)^40,42^ (**Figure S1C**). All cultures reached a TEER peak 8-9 days after reaching cell confluence (−8 to 0 days) indicating the formation of tight junctions. At this point, the cultures were air-lifted and placed into differentiation culture media (21 days). The maturation of the cultures was reflected by mucus production and beating cilia monitored by bright-field microscopy. Cellular composition and appearance of a pseudostratified ciliated columnar epithelium were assessed by immunofluorescence (IF) on frozen tissue sections. All cultures displayed 4 major markers of primary broncho-epithelial differentiation: cytokeratin 5 (Basal cells), SCGB1A1 (club cells), Muc5ac (goblet cells) and acetyl-⍺-tubulin (ciliated cells) (**Figure 1A, Figure S1B and Table S2**). As shown in **Figure 1B**, all donors yielded ALI cultures containing the expected cell subsets, with variable cellular composition between-donors.

**Figure 1.**
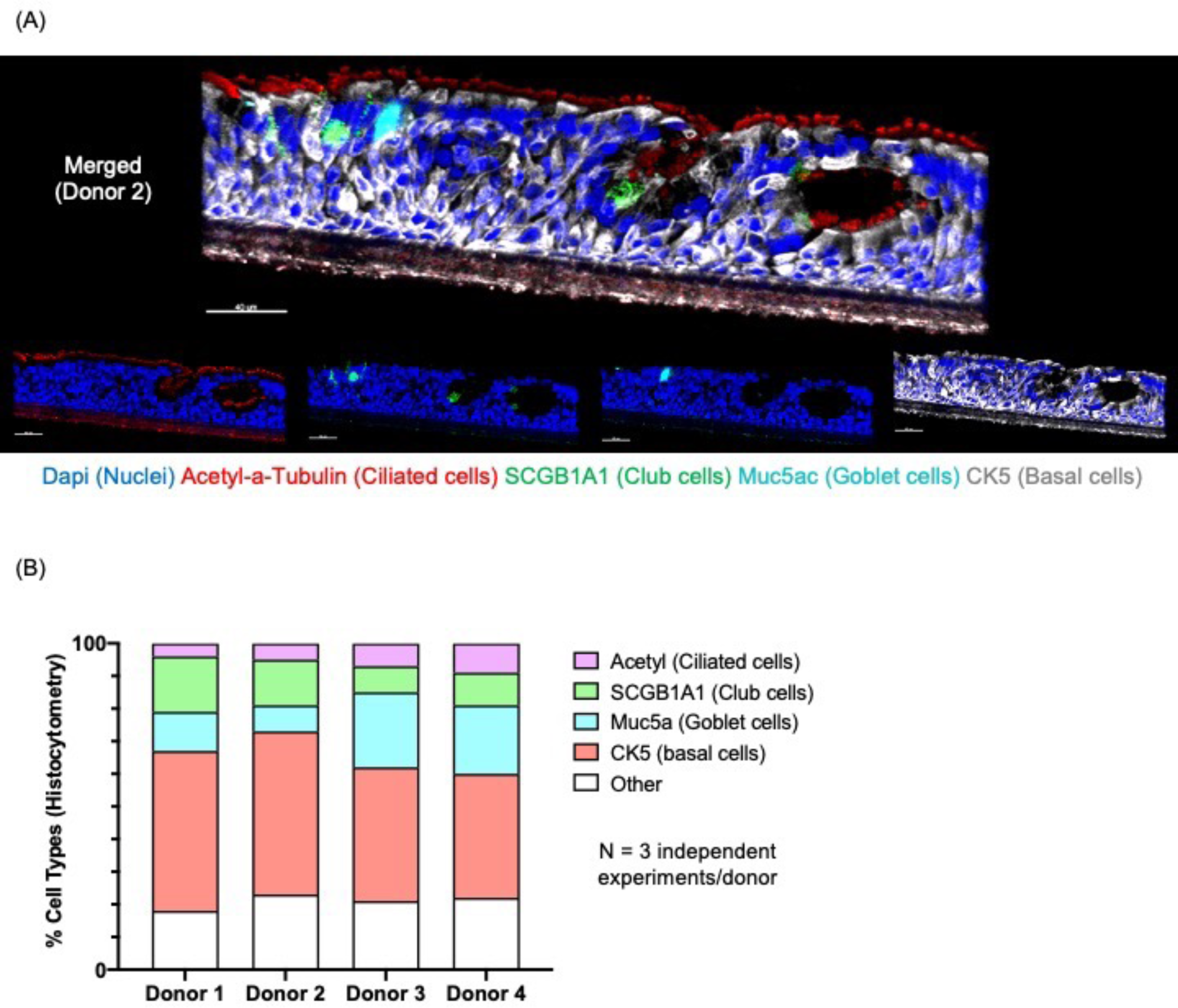
Primary human lung organoid-derived air-liquid interface (ALI) cultures. (A) Representative immunofluorescence staining (IF) of tissue section (8 μm) from differentiated lung organoid-derived ALI cultures of one donor (Donor 2; upper part, merged figure; lower part two-channel figures), showing markers for basal cells (CK5, white), goblet cells (MUC5AC, cyan), club cells (SCGB1A1, green), ciliated cells (acetylated a-tubulin, red) and Nuclei (Dapi, blue). Scale bar 40 μm, in white on the left corner. (B) Quantification of cell types in differentiated primary human lung-derived organoid ALI cultures per donor by histocytometry based on mean staining intensity for the indicated markers from IF scans. Normalized bar graphs represent the percentages of cell markers per donor and were generated using Prism 8 Graph Pad. Data shown are representative of 3 independent experiments per donor with 1 replicate per experiment. See also Figure S1 and Table S1.

### Infection of ALI cultures with SARS-CoV-2 Wuhan-like virus was detected at later time points

To determine the response to virus, we apically applied 10^5^ PFU/well of SARS-CoV-2 Wuhan-like virus in serum free media, (**Figure S2A**) and then harvested wells from 1 to 6 days post infection (DPI). Mock-infected ALI cultures (infection medium without virus) were collected as controls at 6DPI. The presence and magnitude of infection was quantified by multiple assays: 1. IF staining of viral nucleoprotein (NP), followed by microscopy and histocytometry on non-dissociated formaldehyde-fixed tissue sections; 2. Flow cytometry for intracellular NP expression on single cell suspension from dissociated fixed tissue, and 3. Plaque assay on apical supernatant from infected and mock-infected ALI cultures to determine replicating virus titers (**Figure 2 and Figure S2**). Over the six-days post infection, we detected NP staining (indicative of viral replication) in 3 out of four tested donors. Infection was undetectable in ALI cultures from donor 2 in several experiments with Wuhan-like virus. However, infection was detected with delta variant on ALI from the same donor 2 (**Figure S3**). This response could be related to host polymorphism impacting SARS-CoV-2 USA/WA1-2020 virus infectivity^43,44^ and might require further studies. In non-dissociated tissues, NP staining could be detected at the earliest at 3-4DPI, (mean ± SD): 3DPI, 5.1% ± 13 – 4DPI, 7.7 % ± 10, (**Figure 2C and Table S2**). The highest frequency of NP expressing cells was observed at 5-6DPI with 27% ± 26 – 24% ± 18 infected cells, respectively (**Figure 2C, Figure S2C and Table S2**). Similar results were observed using flow cytometry (**Figure 2D, Fig S2B and D, and Table S2**). Viral titer analysis showed the presence of virus in apical washes at 1DPI and a statistically significant increase with time post-infection (mean ± SD) 4.2×10^5^ ± 9.8×10^5^ PFU/ml (4DPI, p-value=0.0301), 9.3×10^5^ ± 1.1×10^6^ PFU/ml (5DPI, p-value=0.0002) and 8.1×10^5^ ± 9.9×10^5^ PFU/ml (6DPI, p-value=0.0014). (**Figure 2E, Figure S2E and Table S2**). Thus, in the absence of other cell types, productive infection of broncho-epithelial cells by the Wuhan-like virus was detected at later time points.

**Figure 2.**
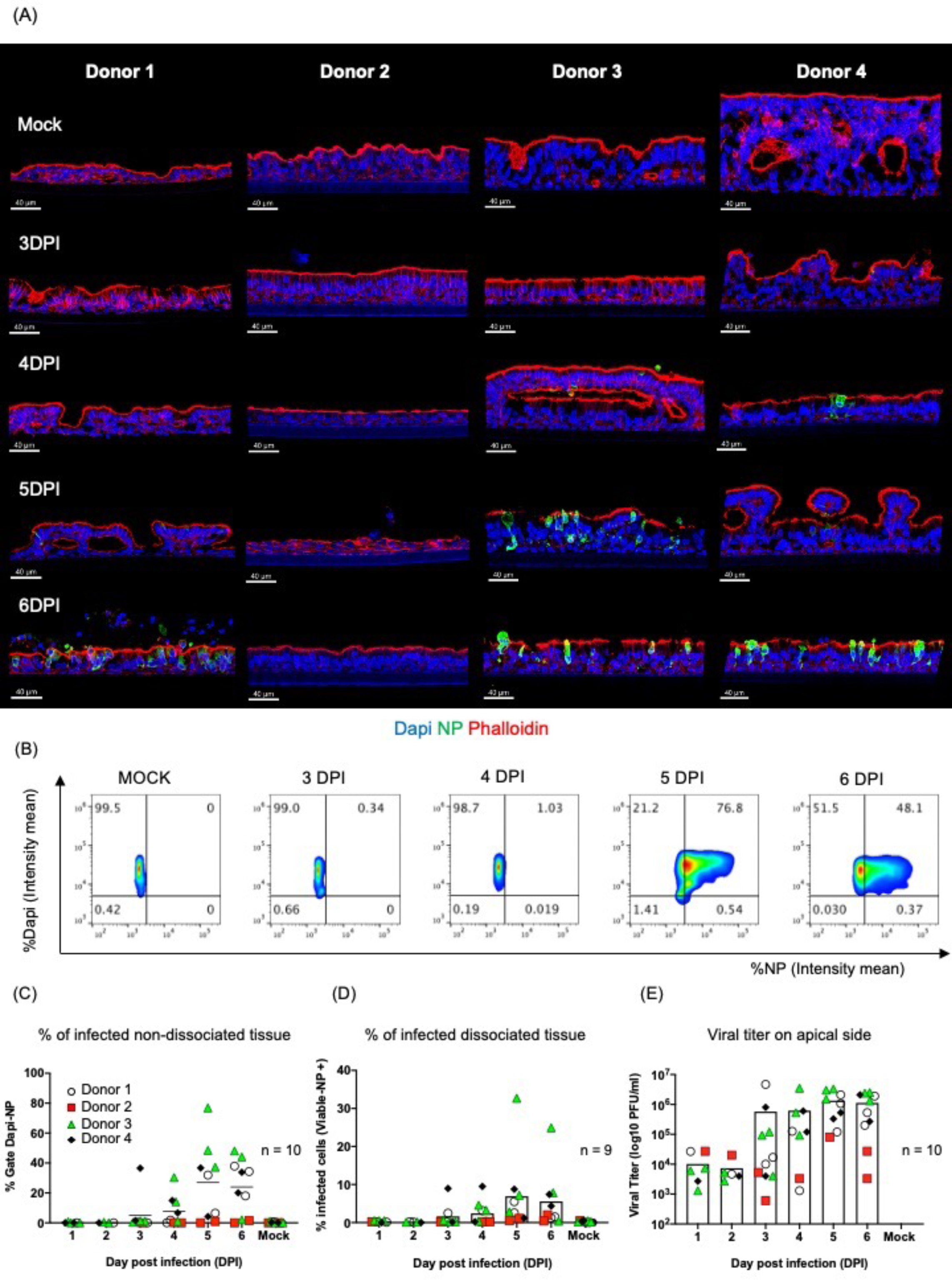
Primary human lung organoid-derived ALI cultures are permissive to SARS-CoV-2 infection (USA/WA1-2020, Wuhan-like virus). (A) Representative images per donor of mock-infected (first line, controls media without virus at 6 days) and infected ALI with SARS-CoV-2 (10^5^PFU) from 3 to 6 days post infection (DPI) stained for nuclei (Dapi), viral nucleoprotein (NP, green) to reveal the effective viral replication and phalloidin (Actin filament, red) to reveal tissue structure. Scale bar 40 μm, in white on the left corner. (B) Representative histocytometry plots on uninfected and infected lung-derived organoids ALI cultures (donor 3) from 3 to 6 DPI. Each dot plot represents the pooling of three consecutive sections, 8 μm thick, showing the mean staining intensity for cell populations positive for Dapi (Y axis) and viral NP (X axis). The NP (for viral infection) channel was generated in Imaris 9.4 using the Channel Arithmetics Xtension prior to running surface creation to identify Dapi-NP cells in images. Statistics were exported for each surface and imported into FlowJo v10.3 for image analysis. (C-E) Quantification of SARS-CoV-2 infection in ALI cultures by different methods: (C) Histocytometry (left scatter plot with bar graph) based on Dapi and NP signal. (D) Flow cytometry (middle scatter plot with bar graph), based on cell viability and NP signal. (E) Viral titer (right scatter plot with bar graph) based on the number of plaques visualized by staining (see material and methods). Data shown are representative of 9-10 independent experiments with at least 2 independent experiments per donor (four donors). Each replicate corresponds to viral titer, or the intensity mean percentage for histocytometry and flow cytometry data, represented by shape and color: Donor 1, white circle; donor 2 red square; donor 3 green triangle and donor 4 black diamond. Bars indicate mean. See also Figure S2 and Table S2.

### SARS-CoV-2 variants of concern display accelerated infection

Using the system described above, we next analyzed the dynamics of ALI infection with Beta (B.1.351), Delta (B.1.617.2) and Omicron (B.1.1.529; BA.1) variants (10^5^ PFU/well) (**Figure 3A**). Based on histocytometry quantification, NP expression in ALI exposed to Beta and Delta variants was already detectable at 3 DPI [(mean ± SD) 10% ± 11 for Beta and 20% ± 23 for Delta, respectively] with peak at 4-5 DPI (**Figure 3B and Table S2**). However, in case of Omicron, NP was detectable at 1DPI (13 % ± 14), with peak at 2DPI (24 % ± 15), then plateau from 3-5DPI before decreasing at 6DPI: 3DPI (23 % ± 9.8); 4DPI (25 % ±16); 5DPI (24 % ± 15) and 6DPI (11 % ± 7.8), (**Figure 3B**, and **Table S2**).This pattern of NP expression was confirmed by flow cytometry (**Figure S3A and Table S2**). As expected, viral particles could be detected starting at 1DPI corresponding to the apical addition of virus with significant increase over time (**Figure S3B and Table S2**). To explore which cells expressed NP, we analyzed infected ALI by IF using markers of the epithelial cell types: cytokeratin 5 (basal cells), SCGB1A1 (club cells), Muc5ac (goblet cells) and acetyl-⍺-tubulin (ciliated cells) as described above. Wuhan-like virus, Beta, Delta and Omicron variants infected predominantly secretory cells (club and goblet cells) and ciliated cells, whereas basal cells were rarely infected (**Figure S4**). Thus, broncho-epithelial cells are permissive to infection with variants. Furthermore, ALI exposed to Delta and Omicron variants show early NP expression suggesting accelerated infectivity.

**Figure 3.**
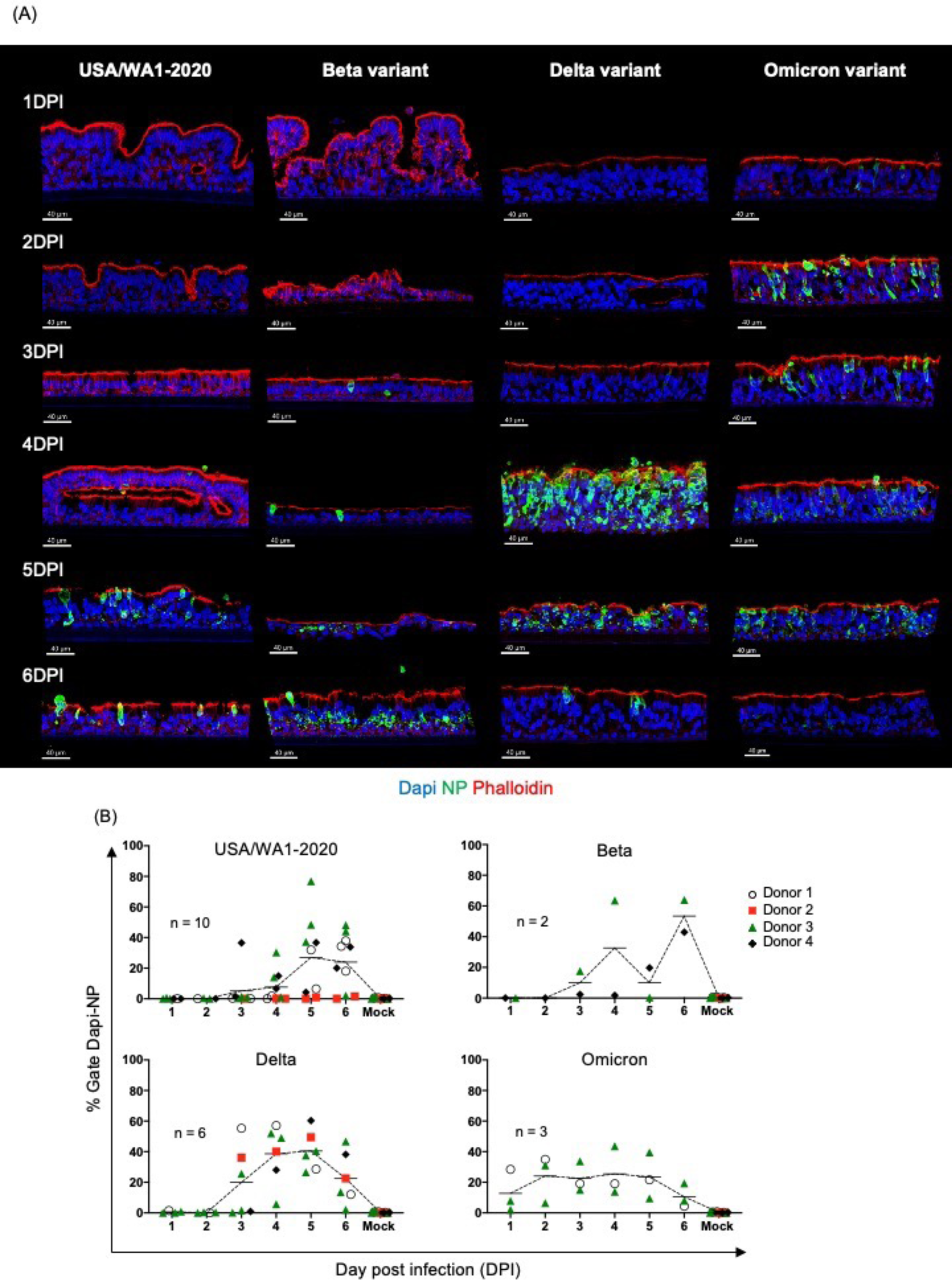
Primary human lung organoid-derived ALI cultures infected with SARS-CoV-2 variants. (A) Representative images (USA/WA1-2020, Delta and Omicron tissue from donor 3 and Beta variant donor 4) per variant of infected ALI with SARS-CoV-2 (10^5^PFU) from 1 to 6 days post infection (DPI) stained for nuclei (Dapi), viral nucleoprotein (NP, green) to reveal the effective viral replication and phalloidin (Actin filament, red) to reveal tissue structure. Scale bar 40 μm, in white on the left corner. (B) Quantification of SARS-CoV-2 infection in ALI cultures by histocytometry per strain: USA/WA1-2020 (top-left scatter line plot graph); Beta (top-right scatter line plot graph); (Delta (bottom-left scatter line plot graph); Omicron (bottom-right scatter line plot graph) based on Dapi and NP signal. Data shown are representatives of at least 2 independent experiments per variant. Each replicate corresponds to the intensity mean percentage represented by shape and color: Donor 1, white circle; donor 2 red square; donor 3 green triangle and donor 4 black diamond. Line indicates mean. See also Figure S3, S4 and Table S2.

### Bi-phasic transcriptional response to SARS-CoV-2 Wuhan-like virus

We next sought to determine the transcriptional response of broncho-epithelial cells to SARS-CoV-2. To start with, we carried out bulk RNA-sequencing (RNASeq) of ALI cultures infected with Wuhan-like virus collected from 1 to 6 DPI and mock-infected control at 6 DPI. Comparative analysis at 6DPI (cut-off: logFC > 1; adjusted p-value < 0.01) yielded a set of 1081 differentially expressed genes (DEGs) up-regulated in ALI exposed with SARS-CoV-2 virus (**Figure 4 and Table S3**). Next, we followed the dynamics and magnitude of changes in expression of these DEGs over time of viral exposure, from 1 to 6 DPI (**Figure 4A**). We identified three clusters: R1 (254 genes) and R2 (150 genes) presenting the most robust transcriptomic changes at 1 DPI, and R3 (677 transcripts) presenting the most robust transcriptomic changes at 5-6 DPI. The early response (R1 and R2) included genes associated with anti-viral status such as *APOBEC3A* and *APOBEC3B*^45,46^; and inflammation^4,47^ such as the IL-1 pathway (*IL1*, *IL1R1, IL1RN, IL17REL, IL32, IL23A, IL36A, IL36G*); *IL-6,* and *TNF* and *NFKB* pathways, (**Figure 4B**). A second group of genes included growth factors (*CSF3*^48^ (granulocytes)); and chemokines involved in the recruitment, differentiation and regulation of immune cells, including *CCL20*^49^ (dendritic cells and lymphocytes), *CXCL1, -3, -6, -8* (neutrophils, granulocytes), *CXCL16* (Macrophages, lymphocytes subsets^50^) and *CXCR4* (regulates cell migration during wound healing). Finally, a group of genes involved in antigen presentation by MHC class I^51^ (*B2M, HLA-A, -B, -C, -E, -G*) and by MHC class II (*HLA-DPB1, HLA-DQB1*); and regulation of adaptive immunity *CD70* and *CD274*. The late response (R3) was dominated by genes associated with viral detection (*DDX58, DHX58, IFIH1, MYD88, TLR3*), IFN type I (*IFNB*), IFN type III (*IFNL1, 2, 3*), and downstream ISGs (**Figure 4C**); and genes encoding for chemokines important for immune cells chemoattraction such as monocytes, T and NK cells including *CX3CL1, CXCL10 and CXCL11*. Overall, these signatures corroborate earlier data on the role of inflammation ^52,53^and IFN response^1,13,54,55^ in SARS-CoV-2 infection and we have now identified a bi-phasic pattern of broncho-epithelial response to Wuhan-like virus.

**Figure 4.**
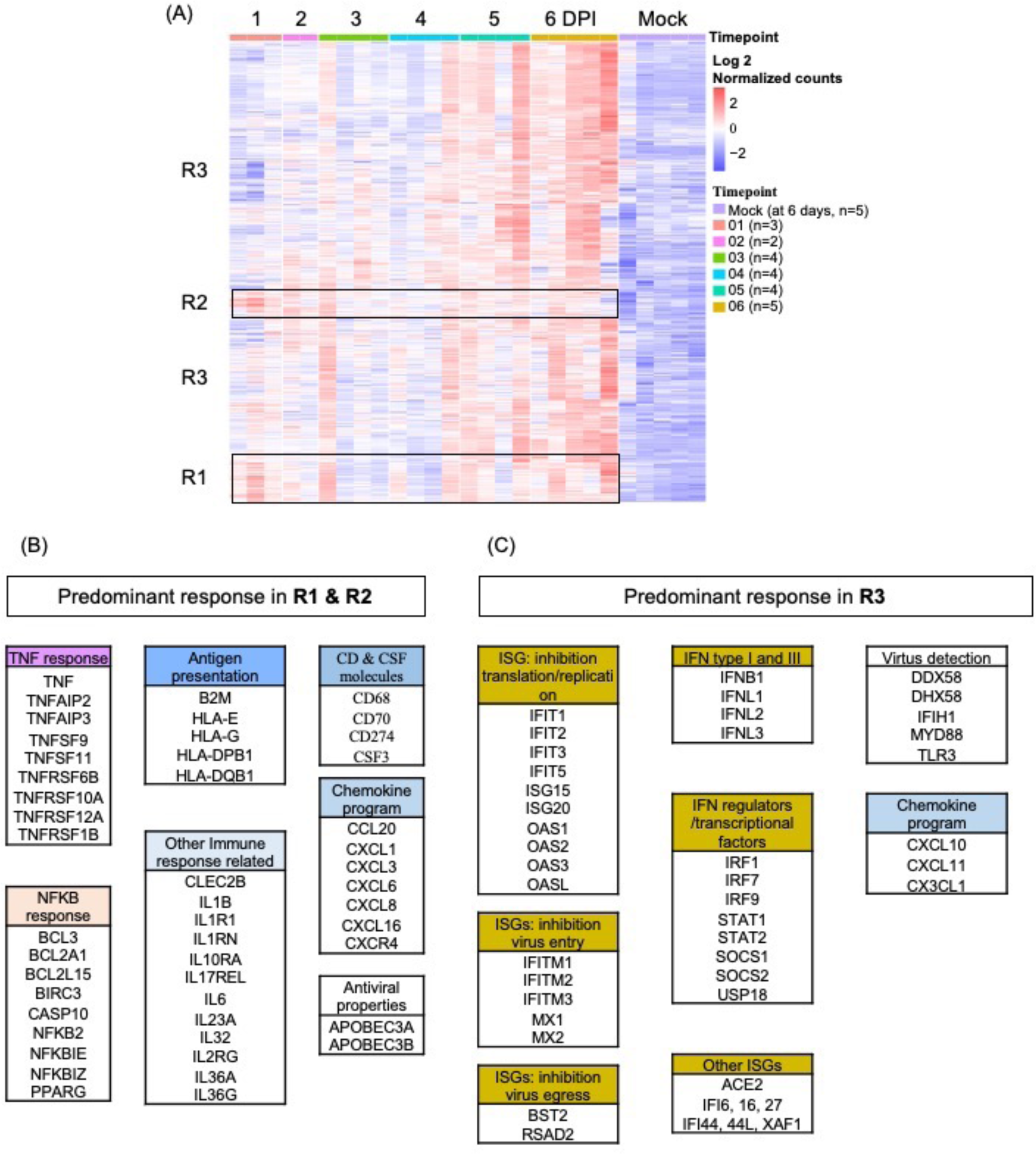
Transcriptional analysis of SARS-CoV-2 (Wuhan-like) infected ALI. (A) Heatmap representing differentially expressed genes (DEG, 1081 transcripts: Log 2 normalized counts) from the comparison 6 DPI vs time-matched mock-infected control (at 6 days) over time in response to SARS-CoV-2. ALI from four donors were infected with SARS-CoV-2 and harvested for sequencing at 1, 2, 3, 4, 5 and 6 dpi and mock-infected samples collected after 6 days (controls media without virus). The sequencing was performed in multiple batches with at least 2 independent experiments at each time-point, the cut-off used: cut-off: logFC > 1; adjusted p-value < 0.01. Rows represent individual transcripts and columns represent individual biological replicates ordered by timepoint. Donor 2 did not show significant response to USA/WA1-2020 infection and was removed from the analysis. Mock-infected samples from one of the batches (donor 1) showed significant differences to the mock-infected samples from different batches and were removed from the analysis as well. Batch effect was removed using SVAseq R package. (B) List of predominant signatures identified in clusters R1 & R2 (early response, robust up-regulation starting at 1DPI). (C) List of predominant signatures identified in cluster R3 (late response, robust up-regulation at 5-6DPI) in response to SARS-CoV-2. See also Table S3.

### Accelerated transcriptional response to SARS-CoV-2 variants

We next sought to evaluate the transcriptional response to different SARS-CoV-2 variants. To this end, we carried out bulk RNA-sequencing (RNASeq) of ALI exposed to different SARS-CoV-2 variants, or mock-infected control, collected from 1 to 6 DPI. To detect genes differentially expressed by tissues exposed to SARS-CoV-2 variants, we used a multi-step analysis, in which Wuhan-like data discussed above alongside data from three variants were incorporated. Briefly, we first compared 6 DPI with the time-matched mock-infected (cut-off: |logFC| > 1; adjusted p − value < 0.01; normalized counts > 10) for each variant independently. Protein coding genes were selected for analysis. We then used DESEQ2 to identify DEGs between variants at each day post-infection, and finally, we overlapped these genes with variant specific DEGs. The combined list had 1608 genes **(Table S4**) whose expression we followed over time (**Figure 5A**). We identified three clusters: one containing genes related to epithelia, cilia and tissue homeostasis programs (750 genes, R4); another enriched in inflammatory and immune programs (313 genes, R5) and finally another enriched in IFN response (545 genes, R6) representing the most robust transcriptomic changes over time and per variant (**Figure 5B-D**). This analysis revealed that variants shared common signatures, with predominant inflammatory and immune response in the earlier time-points, and a predominant IFN response in later time-points. However, differences were found between the variants in the kinetics and magnitude of the transcriptional response (**Table S4)**. To visualize this, we represented each cluster as line plots (each dot plot is the mean of all the log 2 normalized counts of genes in each cluster per time point) in comparison to mock-infected time-matched controls (**Figure 5B-D**). The cluster R4 was enriched for cilia and epithelium maintenance signatures (*KRT1*^56,57^*, POTEJ*^58^*, SERPINA11*^59^*, SERPINA6*^59^*, DAPL1*^60^*, CST6*^61^). Interestingly, this signature was persistently downregulated in response to Delta and Omicron variants. The cluster R5 showed a strong up-regulation of inflammatory and immune signatures for all variants starting at 1 DPI but diverging at 3DPI with an increased inflammation response to Delta variant (**Figure 5C**). Cluster R6 showed progressive and linear increase from 1 to 6 DPI for Wuhan-like, Beta and Delta variants, whereas Omicron follow a parabolic progression until 4 DPI followed by a decrease. All variants induced expression of genes associated with viral detection (*DDX58, DHX58, MYD88, TLR3*), IFN type I (*IFNB, IFNE*), IFN type III (*IFNL1, 3*), downstream interferon-stimulated genes (ISGs) involved in anti-viral activities: inhibition of viral translation/replication (*IFITs, ISG15, ISG20, OASs*); inhibition of virus entry (*IFITMs, MX1, MX2*); inhibition of viral egress (*BST2, RSAD2*); IFN regulators & transcriptional factors (*IRF1, IRF7, STAT1/2, SOCs, USP18*) among others (**Figure 5D**). Overall, our data suggest that SARS-CoV-2 variants show evolution towards accelerated infection which is accompanied by accelerated transcriptional responses in broncho-epithelial cells.

**Figure 5.**
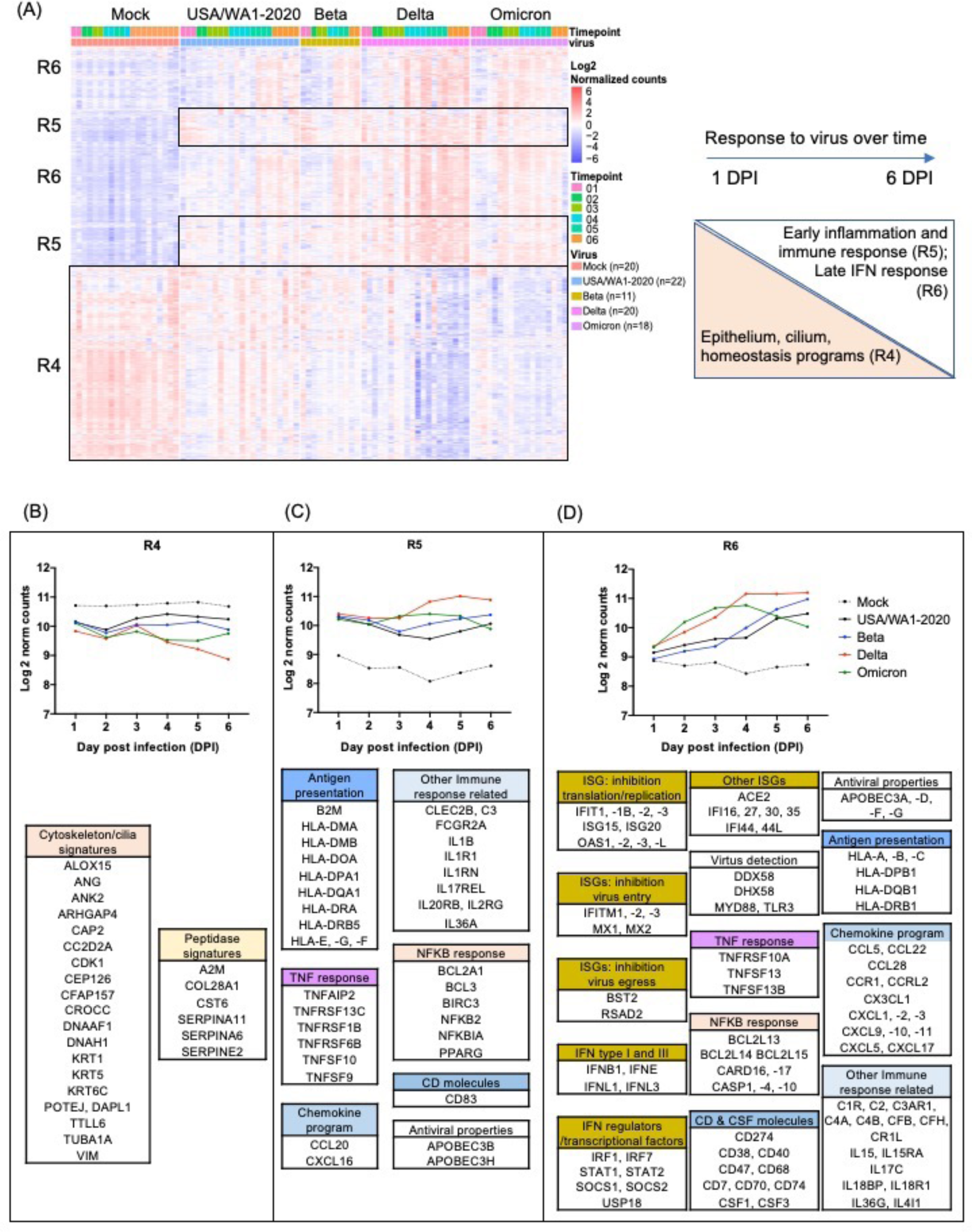
Transcriptional response to SARS-CoV-2 variants. (A) Heatmap representing differentially expressed genes over the time in response to SARS-CoV-2 variants. ALI from four donors were infected with SARS-CoV-2 and harvested for sequencing at 1, 2, 3, 4, 5 and 6 dpi and mock-infected (control media without virus) samples collected from 1 to 6 days as well. The sequencing was performed in multiple batches with at least 2 independent experiments at each time-point, the cut-off used: |logFC| > 1; adjusted p − value < 0.01; normalized counts > 10. Rows represent individual transcripts and columns represent individual biological replicates ordered by timepoints and SARS-CoV-2 variants. Donor 2 did not show significant response to USA/WA1-2020 infection and was removed from the analysis only for Wuhan-like virus. Mock-infected samples from one of the batches (donor 1) showed significant differences to the mock-infected samples from different batches and were removed from the analysis as well. Batch effect was removed using SVAseq R package. (B) Line plots graphs representing the evolution over time of “epithelium, cilium and homeostasis programs”, cluster R4; (C) Early inflammation & immune responses, cluster R5; (D) Late IFN response, cluster R6. Clusters of transcripts in response to SARS-CoV-2 variants, were generated in Graphpad Prism 8. Each dot corresponds to the mean of genes in each cluster per time-point and per strain. See also Table S4.

### Spatial organization of immunomodulatory gene expression in response to SARS-CoV-2

Next, we sought to establish spatial gene expression profiles in ALI cultures exposed to SARS-CoV-2 virus (USA/WA1-2020) using the NanoString GeoMx^®^ digital spatial profiling (DSP) platefrom with the Whole Transcriptome Atlas (WTA) assay^62,63^. First, PFA 4% fixed sections from ALI cultures exposed to USA/WA1-2020 (from 1 to 6 DPI) or mock-infected (at 6 day) were labelled with antibodies targeting NP and Spike viral proteins, to reveal infected cells, and with CK5 antibody to identify and visualize basal cells (**Figure 6A-B**). Second, *in situ* hybridization probes targeting 18,676 total human transcripts were used, each probe tagged with a unique molecular identifier (UMI) for next-generation sequencing. Guided by the antibody staining, we selected at least six regions of interest (ROI) per condition, total 78 ROIs (**Table S5**): apical cells CK5^-^ and basal CK5^+^ cells from mock-infected controls and from SARS-CoV-2 infected ALI. Probe UMI counts from each ROI were obtained to provide spatial transcriptomes (**Table S6**). Sequencing quality was analyzed to ensure sensitivity of low expressors (**Figure S5A**). Data were normalized to the third quartile (Q3) to account for differences in cellularity, and ROI Size (**Figure S5B**). None of the 78 ROIs analyzed were below the 50% warning (18,676 total targets); 17,266 genes normalized by 3rd quartile, were expressed above the limit of quantification (LOQ) in at least 95% ROIs (**Figure S5C-D**).

**Figure 6.**
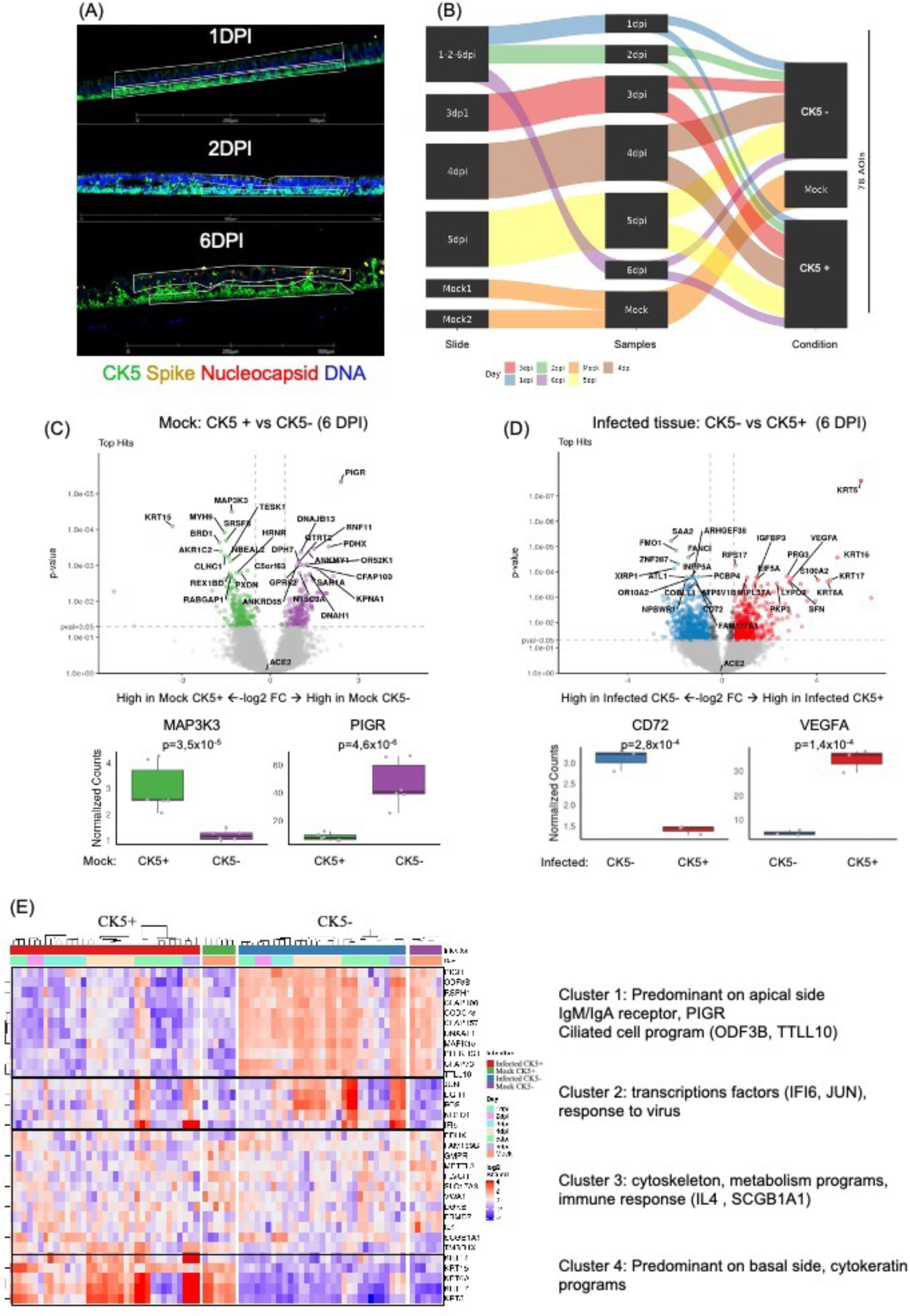
GeoMX digital spatial profiling of SARS-CoV-2 infected ALI. (A) Representative images of infected ALI sections with SARS-CoV-2 (10^5^PFU, from up to bottom), 1 DPI, 2 DPI, 6DPI stained for nuclei (Dapi), viral nucleoprotein (red), spike viral protein (yellow) and cytokeratin 5 (CK5). The thin white polygons represent the ROI strategy selection for the apical cytokeratin 5^-^ cells (CK5^-^) vs basal side CK5^+^. Scale bar 500 μm for 1 and 6 DPI; and 1 mm for 2 DPI (white). (B) Schematic experimental design for ROI selection per slide, time-point (1 to 6 DPI) condition (infected or mock-infected control at 6 days) and apical or CK5^+^, a total of 78 ROI were selected (C-D) Volcano plots visualizing significant differentially expressed genes between two groups of samples (X axis, Log2 FC and Y axis, p-value <0.05), boxplot represents relevant top hit up-regulated in each region (CK5^+^ or CK5^-^). (C) Comparison Mock CK5^+^ vs Mock CK5^-^ (6DPI). (D) Comparison Infected CK5^-^ vs Infected CK5^+^ 6DPI). (F) Heatmap representing the 33 tops differentially expressed genes (DEG) comparing conditions per location-infection status, then per timepoint (1 to 6 DPI). The sequencing was performed in 2 batches with at least 3 replicates per condition from two representative experiment (donor 3 for slides #1, #2, #4 and #5; on slide #4, 2 sections from donor 4 and one from donor 3). See also Figure S5, Table S5 and Table S6.

We first compared the CK5^+^ (basal cells) to CK5^-^ (apical cells) cells in mock-infected controls to establish predominant signatures characterizing these regions in non-infected tissue. Basal CK5^+^ region was enriched in transcripts related to cellular cytoskeleton compounds ensuring the basal layer maintenance (*KRT15, MYH9, BRD1*) and cell proliferation (*MAP3K3*)^64^. The apical CK5^-^ region was enriched in transcripts related to ciliated cells, as well as *PIGR* which is a receptor allowing the basal to apical transcytosis of IgA and IgG (**Figure 6C**). To determine the “spatial” response to SARS-CoV-2 Wuhan-like virus infection, we compared infected CK5^-^ to CK5^+^ cells at 6 DPI (**Figure 6D**). Basal CK5^+^ infected tissue, was enriched in transcripts related to cell damage/repair (*VEGFA*^65^*, IGFBP3*^66^*, EIF5a, S100A2*^67^*, PKP3, LYPD3, SFN*), cytokeratins (*KRT5, KRT6A, KRT16, KRT17*) possibly related to basal cell differentiation/proliferation properties for maintenance/repair of airway epithelium^68–72^, (**Figure 6D-E**). Apical CK5^-^ infected tissue, was enriched in transcripts related to actin filaments/cilia (*ATL1, COBLL1, OR10A2, PCBP4, INPP5A, XIRP1*), regulatory factors (*ARHGEF38, ZNF287, FANCI, FMOI, NPBWR1*), as well as *CD72* (invariant chain) and inflammation (*SAA2*) (**Figure 6D**).

The DSP analysis suggested spatial organization of airway epithelial response to virus. RNAseq analysis revealed up-regulation of *CSF3* and *CCL20*, critical signals for differentiation and regulation of immune cells, starting at 1DPI upon SARS-CoV-2 exposure. Therefore, we analyzed the localization of CSF3 and CCL20 protein expression by IF staining on fixed tissue sections of ALI over time following SARS-CoV-2 exposure. Tissue staining revealed a consistent pattern of CSF3 protein expression in basal layer of cells (**Figure 7A**) and this staining localization was common across ALIs exposed to different SARS-CoV-2 variants (**Figure S6A**). Furthermore, in normal human lung, CSF3 staining was localized predominantly in the basal layer of lung epithelium as delineated by expression of CK5 and the phalloidin which is bright and strong in epithelium tight junctions/cilia (binds to actin filament), (**Figure 7B and Figure S6B**). In addition, on a consecutive tissue section of normal human lung, we stained for immune cells markers such as CD11b and CD14 and found that in the same lung epithelium area, CSF3 was localized in proximity of CD11b+ myeloid cells (**Figure 7C and Figure S6C**). CCL20 protein staining was present at low levels throughout the ALI tissue in mock-infected condition (**Figure 7A**). However, upon SARS-CoV-2 exposure its expression increased and localized to apical side (**Figure 7A**). This CCL20 expression pattern was common across ALIs exposed to different SARS-CoV-2 variants (**Figure S6A**). In normal human lung, at steady state, CCL20 was present throughout the tissue (**Figure 7B-C and Figure S6B-C**), reflecting the staining pattern visualized in ALI mock-infected condition.

**Figure 7.**
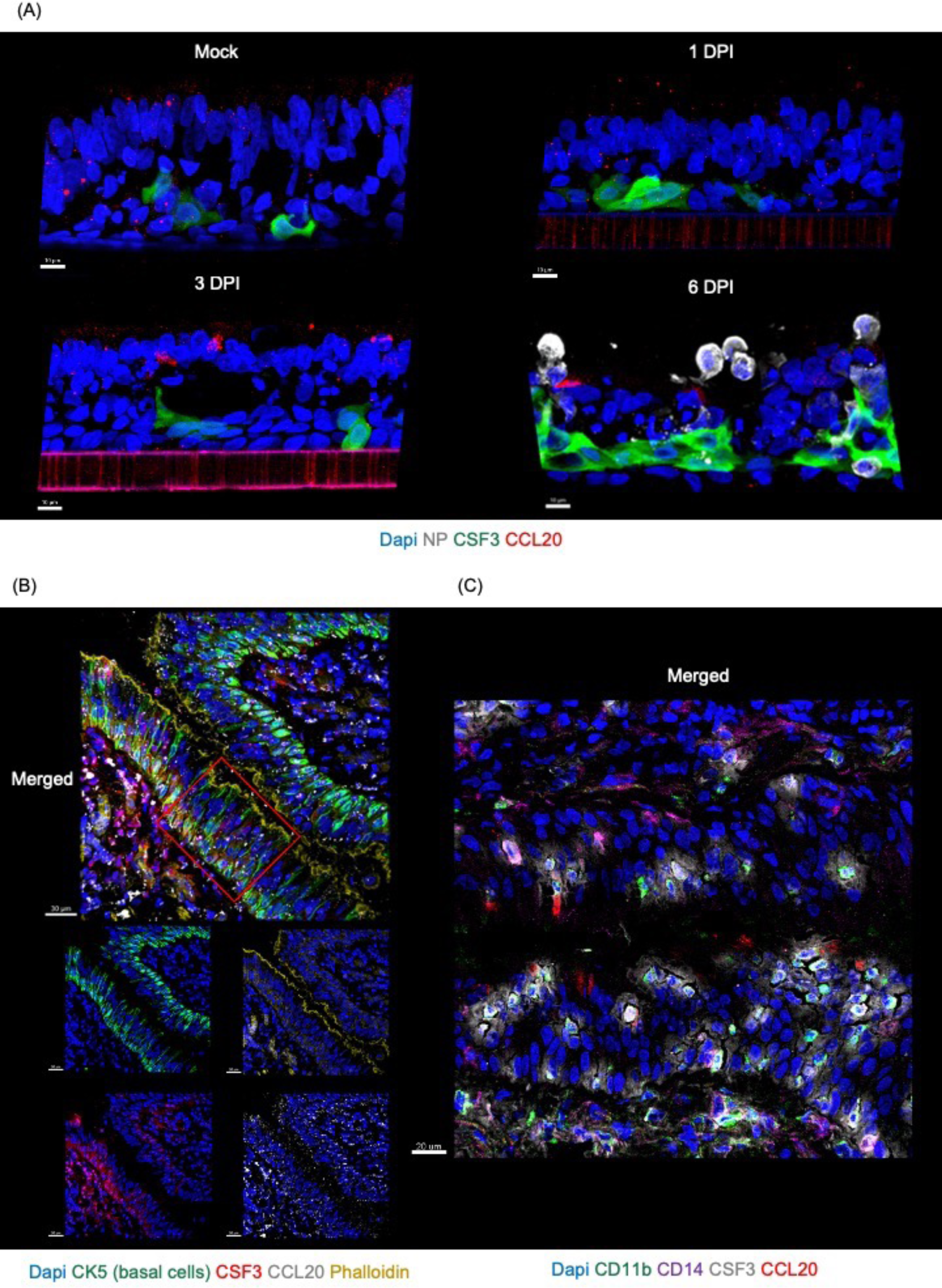
CSF3 and CCL20 expression in ALI exposed to SARS-CoV-2 USA-WA1/2020 and in human lung epithelium. (A) Representative images of ALI (donor 3) mock-infected at 6 days and ALI infected with SARS-CoV-2 USA/WA1-2020 (10^5^PFU) at 1, 3 and 6 days-post infection (DPI) stained for nuclei (Dapi, blue), viral NP (white) to reveal the effective viral replication, CSF3 (green) and CCL20 (red). Scale bars 10 μm, in white on the left corner. (B) Representative images of human lung epithelium (Donor 1) stained for nuclei (Dapi, blue), CK5 (green), phalloidin (Actin filament, dark yellow), CSF3 (red) and CCL20 (white). Scale bars 30 μm, in white on the left corner. (C) Representative zoomed image of human lung epithelium (Donor 1) stained for nuclei (Dapi, blue), CD11b (green), CD14 (magenta), CSF3 (white) and CCL20 (red). Scale bars 20 μm in white on the left corner. See also Figure S6.

## Discussion

Using human broncho-epithelial air-liquid-interface (ALI) tissue cultures which resemble the conditions of human airway epithelia, here we show that SARS-CoV-2 virus and its variants trigger epithelial cell-type specific transcriptional response that is modulated over time to engage different arms of the immune system. First, we find that infection defined by the presence of NP staining, was a late event in SARS-CoV-2 Wuhan-like virus and Beta variant, whereas it was accelerated in the cases of Delta and Omicron variants. This difference between variants, could be due to the accumulated mutations in Delta and Omicron modifying their ability to form syncytia or entry into host cells. It has been reported in previous studies, that Delta had increased capacity to infect cells expressing low levels of ACE2 and form syncytia by comparison to other variants and presented increased capacity for epithelial damage^32,35,73^. Interestingly, Omicron displays a lesser capacity to form syncytia, but has enhanced replication in upper airways possibly due to an endocytic pathway mechanism increasing cell entry^38,74,75^. Variants did not appear to differ in term of cell subtype infection in the ALI model and ciliated and secretory cells were generally infected by all variants, while basal cells were found infected rarely thus corroborating previous studies^35,76^. Overall, the faster kinetics of infection of human ALI cultures SARS-CoV-2 Delta and Omicron variants as compared to previous variants will be consistent with increased adaptation of the more recent variants to infect human respiratory epithelium.

Dynamics of transcriptional response followed infection kinetics. To this end, all variants triggered transcriptional inflammatory, immune response and IFN signatures as revealed by bulk RNAseq. However, differences in the quality and kinetics of transcriptional response to variants were found with SARS-CoV-2 Wuhan-like virus, triggering early inflammatory signature at day 1 preceding a late type I/III IFN response (5-6 DPI), consistent with the timeline of NP protein expression. The response to SARS-CoV-2 Wuhan-like virus was spatially organized as illustrated by GeoMX digital spatial profiling of basal (cytokeratin 5 [CK5^+^] cells) vs apical side (CK5^-^ cells) revealing the presence of cytokeratin signatures, proliferative and regenerative programs in CK5^+^ cells; whereas in CK5^-^ cells, we observed ciliated cells and *CD72* (invariant chain)^77^ programs including common signatures such as MHC-class I (*HLA-A/B/C*) programs and IFN response signature in response to virus. This suggests a potential contribution to antigen specific T cell activation. The rare infection of basal cells, as mentioned above, seemed to respond to viral presence by turning on differentiation markers, consistent with a tissue repair response. While apical cells presented up-regulation of molecules involved in IgA/IgM transcytosis with *PIGR* response and actin filaments/ciliated cell programs, perhaps ready to contribute to viral clearance. Spatial organization was further illustrated at the protein level by CSF3 expression in basal cells and CCL20 expression in apical cells. CSF3 is an important survival and proliferation factor for neutrophils and, along with CXCL6, it could also trigger their recruitment to the damaged area^78^. From single cell data, *CSF3* has been associated to a subset of goblet cells^69^ in the airway epithelium. In ALI cultures models, it has been reported that CSF3, CXCL6, CXCL16 (protein level) were part of the RSV pathological signature^79^, similar patterns were observed in SARS-CoV-2 studies that have reported an up-regulation of *CSF3*^48^. Thus, CSF3 has been suggested as a potential therapeutic target, in which early blocking could help to reduce the hyper-inflammation mediated by exaggerated neutrophilic response^80^. Translocation of CCL20 to the apical side upon viral exposure might help attract neutrophils from basal layer to ciliated cells^49^. Indeed, while CCL20 has an impact on a broad range of cells of the adaptive immune response, it can also attract neutrophils^81^. Finally, *CCL20* and *CSF3* were also induced by the variants used in this study. There were, however, some differences in protein expression. For example, CSF3 was induced as early as 1 DPI but was expressed more highly at later time points and persistent expression even after infection subsided as was the case for Omicron variant. CCL20 was induced in all variants and its expression was sustained in Delta and Omicron.

Overall, we identified three transcriptional clusters present in ALI exposed to SARS-CoV-2 variants. One cluster, R4, enriched in genes involved in ciliated and cytoskeleton homeostasis/organizational programs, which is consistent with the impact of viral infection, clearance and tissue repair^76,82,83^. Clusters R5 and R6 consisted of inflammatory, immune response and IFN signatures^54,84,85^. Interestingly, the kinetic and magnitudes were variant-related, and we found that Wuhan-like and Beta were similar to each other while Delta and Omicron diverged, especially at later time points. The cluster R5 was enriched in inflammation and immune signatures including *TNF*, *HLA* (class I and II), *NFKB*, *IL17REL* and *IL1*. This early response could be consistent with a TLR response characterized for a more skewed NFK-B^86^ response, perhaps RNA/TLR3 sensing triggered by extracellular viral products. The cluster R6 contained among others *IL17C* corroborating the role of IL17 pathway in inflammation^87^ as well as tissue injury/repair ^88^. In terms of IFNs an IFN-signature, Wuhan-like and Beta presented a delayed pattern (3-4DPI), Delta (2-3DPI) and Omicron starting at 1 DPI with IFN type I/III response^54,89^. This delayed pattern of expression was consistent with kinetics of NP expression, reflecting intracellular replication consistent with sensing by RNA cytoplasmic receptors such as MDA5^90^, that are known to be more skewed towards IFN and ISG induction through engagement of IRF3^86^. Thus, IFNs transcript abundance was most pronounced at the time of the highest NP expression suggesting direct correlation between viral replication and IFN response. It is somewhat surprising as it suggests that virus is not sufficiently sensed by epithelial cells at the early time points, which could trigger a potentially protective pathway. While it remains to be determined, possibly accessory cells are needed for the first wave of type I/III IFNs. Thus, SARS-CoV-2 virus triggers a cell-type specific transcriptional response in broncho-alveolar epithelium that is modulated over time to engage different arms of immune response.

## Acknowledgments

We thank our tissue donors. The Microscopy, Single Cell Biology, Genome Technology, and CTRS Scientific Services of The Jackson Laboratory. We thank Randy Albrecht for support with the BSL3 facility and procedures at the Icahn School of Medicine at Mount Sinai (ISMMS). We thank Stefanie Roth for support with sample procurement from Hartford Hospital. The graphical abstract and Figure S2A were partly generated using Servier Medical Art, provided by Servier, licensed under a Creative Commons Attribution 3.0 unported license.

## Author contributions

D.C.C. and S.J. performed the research, analyzed, discussed, interpreted data and literature. D.C.C. drafted the manuscript. M.Y. and J.L.: computational, statistical analysis and manuscript reading. J.M., M.C. (Megan Callender), M.C.: experiment performance, manuscript reading and data analysis. A.C., J.G.-B.D.: experiment performance. T.-C.W: experiment performance, library preparations and manuscript reading. F.M.: tissue sample processing. P.Y., A.S., J.K. G.A.: contributed to the research. B.H., S.E.C., K.G. and E.A.: GeoMx experiment performance, computational data analysis and manuscript reading. D.C., A.G.-S, M.S., A.W. supervised, verified data analysis, and contributed to manuscript critical reading. K.P.: designed the study, supervised, discussed the results, and co-wrote the manuscript.

## Declaration of interest

The A.G.-S. laboratory has received research support from GSK, Pfizer, Senhwa Biosciences, Kenall Manufacturing, Blade Therapeutics, Avimex, Johnson & Johnson, Dynavax, 7Hills Pharma, Pharmamar, ImmunityBio, Accurius, Nanocomposix, Hexamer, N-fold LLC, Model Medicines, Atea Pharma, Applied Biological Laboratories and Merck, outside of the reported work. A.G.-S. has consulting agreements for the following companies involving cash and/or stock: Castlevax, Amovir, Vivaldi Biosciences, Contrafect, 7Hills Pharma, Avimex, Pagoda, Accurius, Esperovax, Farmak, Applied Biological Laboratories, Pharmamar, CureLab Oncology, CureLab Veterinary, Synairgen, Paratus and Pfizer, outside of the reported work. A.G.-S. has been an invited speaker in meeting events organized by Seqirus, Janssen, Abbott and Astrazeneca. A.G.-S. is inventor on patents and patent applications on the use of antivirals and vaccines for the treatment and prevention of virus infections and cancer, owned by the Icahn School of Medicine at Mount Sinai, New York, outside of the reported work.

The M.S. laboratory has received unrelated funding support in sponsored research agreements from Phio Pharmaceuticals, 7Hills Pharma, ArgenX and Moderna.

S.E.C. declares being an employee and stockholder at NanoString Technologies.

KP is a stockholder in CueBiopharma and Guardian Bio, scientific advisor to Cue Biopharma and Guardian Bio and co-founder of Guardian Bio. K.P. declares unrelated funding support from Guardian Bio (current) and MERCK (past). All additional authors declare no competing interests.

## Inclusion and Diversity

We support inclusive, diverse, and equitable conduct of research.

## Material and Methods

### Key resources table

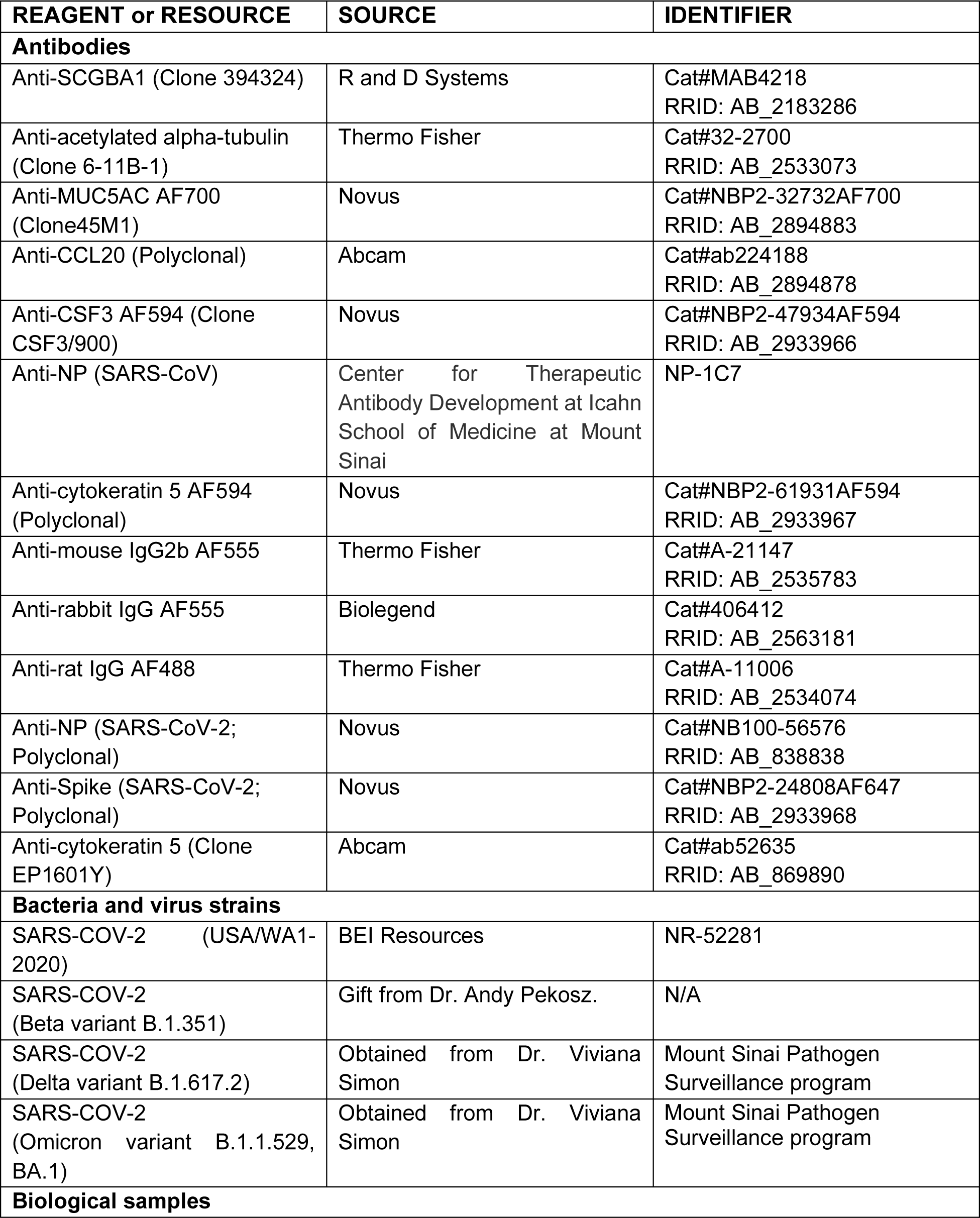

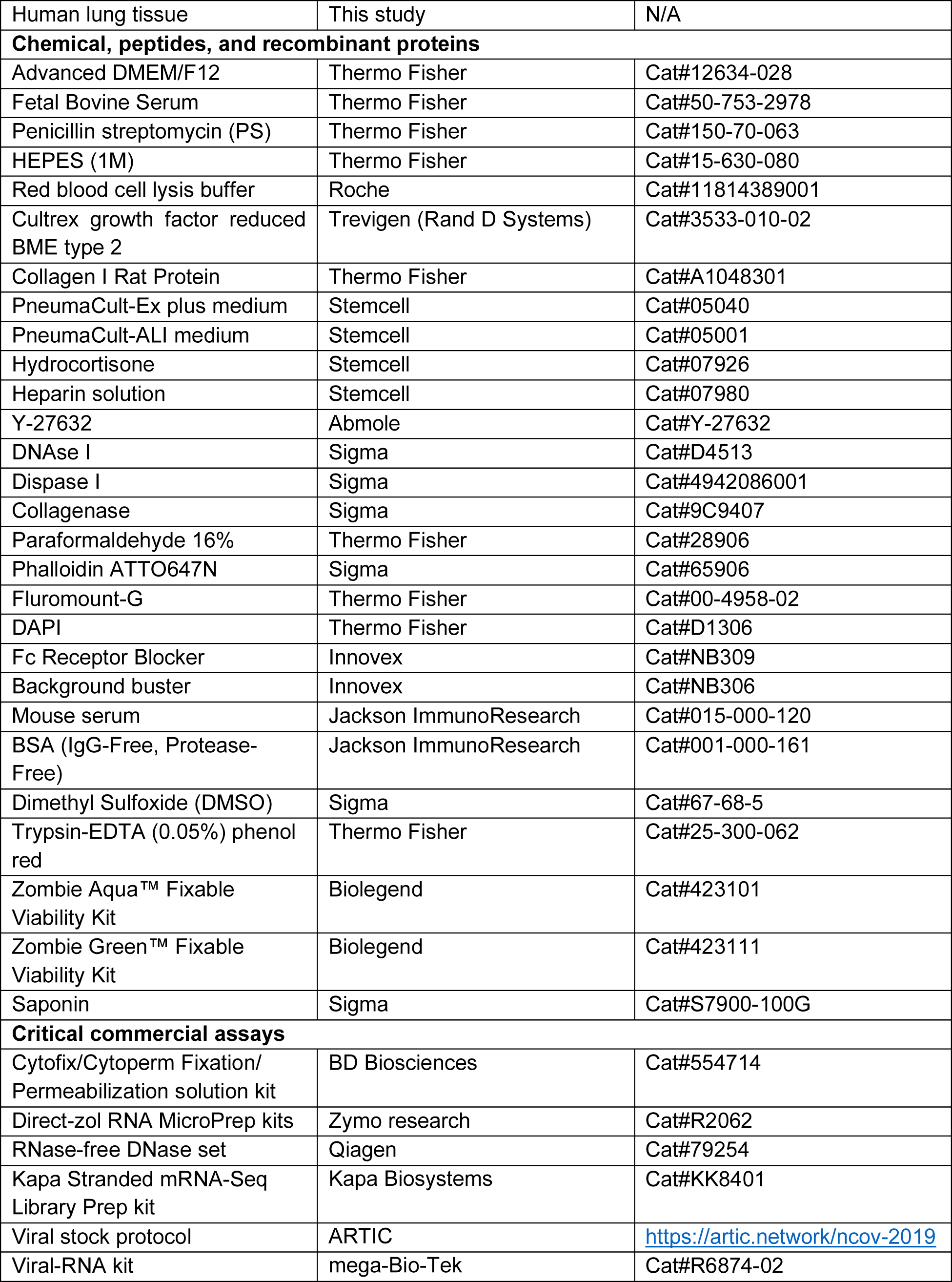

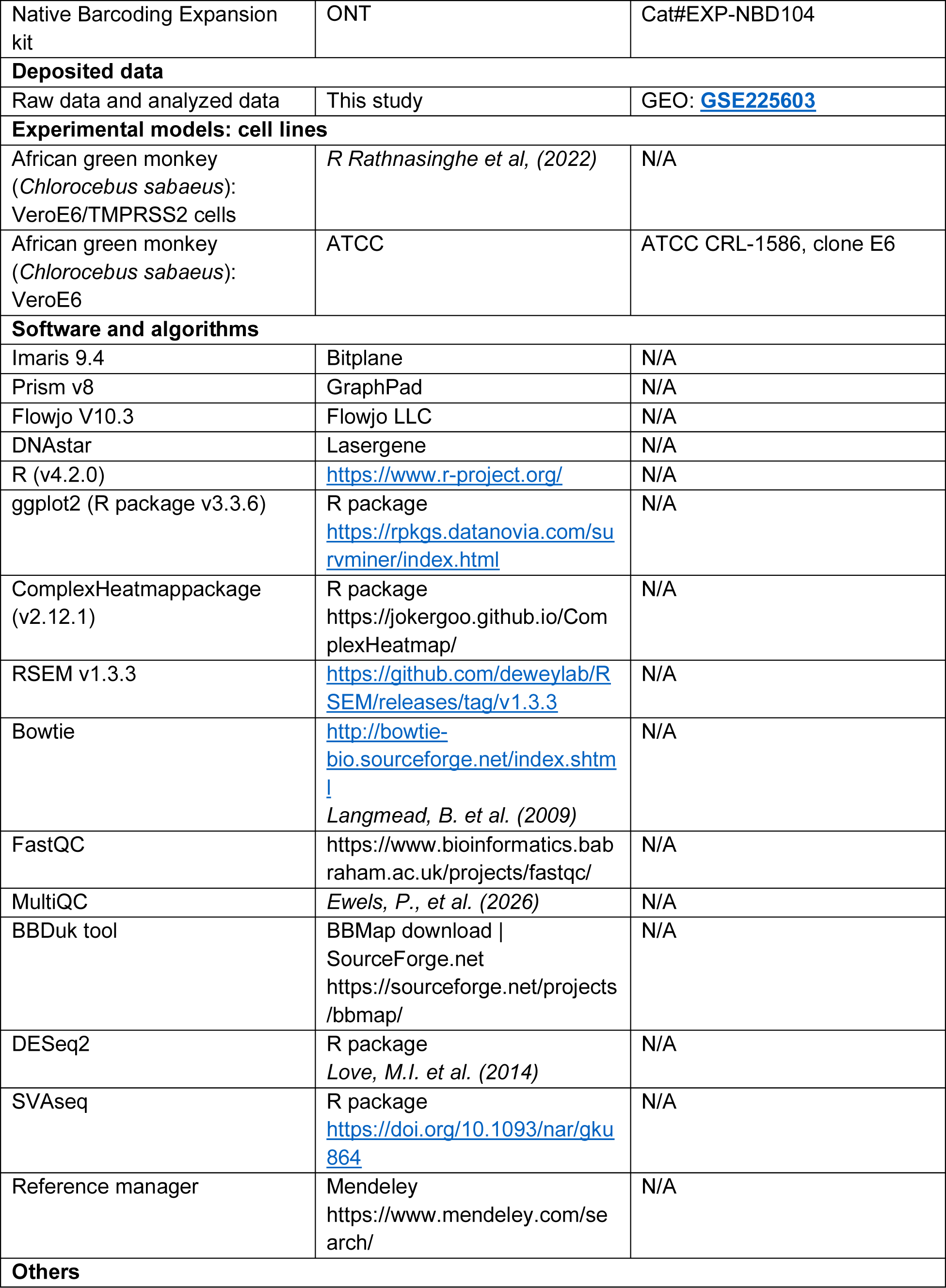

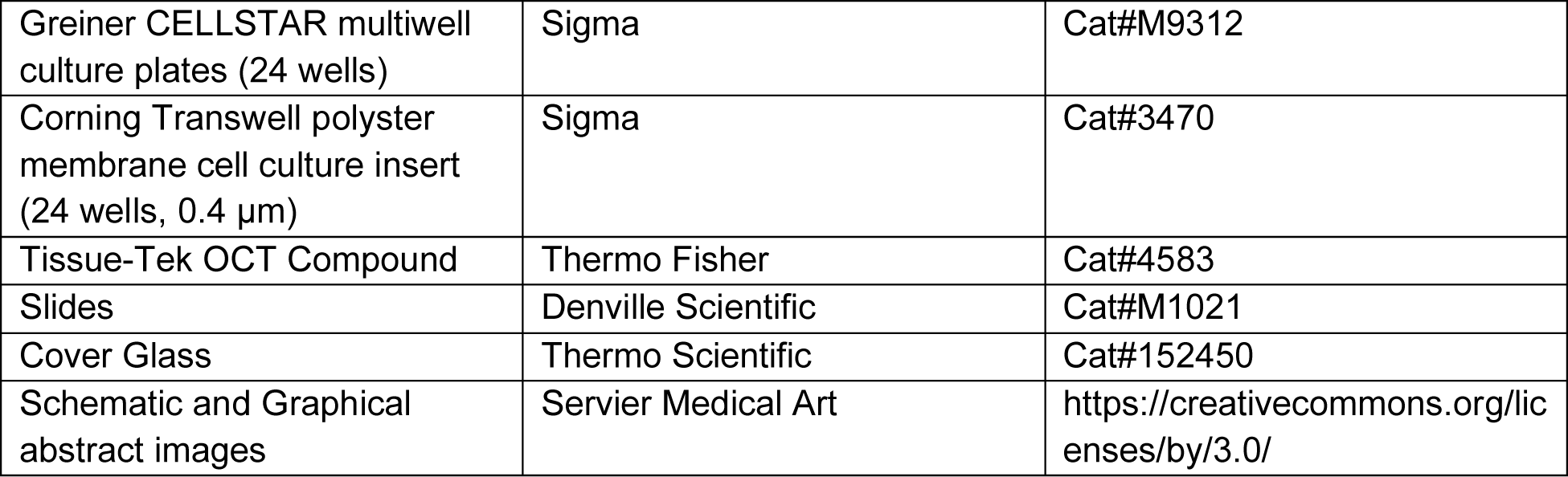

## Resource availability

**Materials availability:**

This study did not generate new unique reagents.

### Data and code availability

- All sequencing data generated for this study has been deposited to GEO and is publicly available: **GSE225603**
- This paper does not report original code.
- Any additional information required to reanalyze the data reported in this work paper is available from the lead contact upon request.

## Experimental model and subject details

### Primary lung-organoid generation

#### 1. Procurement of human material and informed consent

De-identified human lung tissues were obtained from NDRI (Project: RPAK1 01), in compliance with relevant American laws and institutional NIH/NIAID guidelines. Study procedures were performed in the context of U19AI142733 grant at the Jackson Laboratory. The overall goal is to elucidate the innate immune networks that shape adaptive immune responses to respiratory viral infections in the human lung. Donors’ demographics are detailed in Table S1.

#### 2. Tissue processing and organoid culture

To generate primary lung organoids, we adopted Sachs et al.^41^ protocol. Briefly, cryopreserved lung tissue (FBS 10% DMSO) was washed with Advanced DMEM/F12 media containing 1x Glutamax, 10mM Hepes and antibiotics (AdDF++), and digested in 10 ml of complete media for organoids (AO), containing 2 mg/ml of collagenase (Sigma) on an orbital shaker at 37°C for 1–2 h. The digested tissue suspension was sequentially sheared using sterile slides and strained over a filter (100 μm) with 10ml AdDF++ media until obtain a single cell suspension. After inactivation with 2% FBS, red blood cell lysis buffer (Roche,) treatment could be performed if needed. Then, cell counting, centrifugation (400 rcf), and resuspend the lung cell pellet in 10mg/ml Matrigel (Cultrex growth factor reduced BME type 2; Trevigen). Embedded Matrigel cells were dispensed as follow: 3.10^5^ cells in a 40μl drops per well in a pre-warmed 24-well plate (Greiner, M9312). Those organoid cultures were maintained over 4-7 passages until obtaining a stock of pure primary lung-organoid derived epithelial cells.

## Method details

### Lung organoid-derived air-liquid interface (ALI) cultures

To generate air-liquid-interface (ALI) cultures, lung-organoid derived epithelial cells were plated on 0.03mg/ml collagen I rat protein (Gibco™ Collagen I Rat Protein, Tail) coated inserts with a pore size of 0.4 μm (Corning 3470) at 3.10^4^ cells/ml per insert into 24-well culture plate. Once the cells were seeded, the procedure of lung-organoid derived ALI culture generation included the three following steps according to manufacturer’s instructions (STEMCELL Technologies, Cambridge, Massachusetts, USA):

1. Maintenance in Pneumacult-Ex Plus Medium containing the ROCK inhibitor Y-27632 until confluence (8-10 days). Fresh medium, 100 μl (apical chamber) and 500 μl (basal chamber) changed every 2 to 3 days.
2. First part of ALI differentiation in Pneumacult-ALI Medium (STEMCELL) containing the ROCK inhibitor Y-27632 (5-7 days). Fresh medium, 100 μl (apical chamber) and 500 μl (basal chamber) changed every 2 to 3 days. At this step, TEER started to be measured every 2-3 days until full differentiation as described previously^40,91^.
3. Second part of ALI differentiation in Pneumacult-ALI Medium (STEMCELL) without ROCK inhibitor Y-27632 (∼28 days) and removal of the media at apical side (air-lift). Fresh medium, 500 μl (basal chamber) changed every 2 to 3 days. The air-lift step is performed once the TEER reach values 500 (ohms/cm^2^) <, indicating healthy confluent layer. Once airlifted, cultures are checked under microscope every 2-3 days for cilia beating appearing in general by 2 weeks and mucus should be present from 2-4 weeks.

All experiments were performed using lung organoid-derived epithelial cells from 4 different donors, for each donor a single lot and single aliquots were used to avoid batch variation.

### Viruses and cells

All experiments involving handling of live SARS-CoV-2 isolates were performed in a biosafety level (BSL) 3 facility at Icahn School of Medicine at Mount Sinai, New York. SARS-CoV-2 (USA/WA1-2020, Wuhan-like), Beta (B.1.351), Delta (B.1.617.2) and Omicron (B.1.1.529, BA.1) variants were obtained through our collaboration with Adolfo García-Sastre’s lab. Infectious stocks were expanded on VeroE6 for USA/WA1-2020 or VeroE6-TMPRSS2 cells for all other SARS-CoV-2 variants. The supernatants were centrifuged and passed through a 0.45-μm filter to get rid of cells. Viral titers were determined by plaque assay on VeroE6 for USA/WA1-2020 or VeroE6-TMPRSS2 cells for all other SARS-CoV-2 variants. All the viral stocks were sequenced to confirm genomic integrity according to the previously described ARTIC protocol (https://artic.network/ncov-2019)^92^. Aliquots were stored in a secured −80 °C freezer until use.

### Virus infection of primary lung-organoid derived ALI cultures

Prior to infection, the mucus from the apical side of primary lung-organoid derived ALI was washed three times with PBS. Viruses were diluted in 1X PBS for a final concentration of 10^5^PFU per ALI for SARS-CoV-2 infections and spread out on apical side. To collect viral supernatant, 150μl of 1X PBS was added on apical side and incubated for 15 min at 37°C.

### Quantification of infectious virus

For viral titration, plaque assays were performed with a 10-fold serial dilution of viral supernatant in 1X PBS containing 1% Bovine serum were overlaid on pre-seeded VeroE6 cell (for USA/WA1-2020) or Vero-E6 TMPRSS2 (for Beta, Delta and Omicron) monolayers and absorbed incubated at 37°C for 1 hour with gentle shaking every 5 min. After incubation, 2% Oxoid agarose mixed with 2X MEM supplemented with a final 0.3% FBS was overlaid, and the cells were incubated for 72h at 37°C. Plaques were visualized using immune staining with SARS-CoV-2 NP antibody (1C7,). Briefly, followed by formaldehyde fixation, the cells were incubated with a mouse anti-SARS-CoV-2 NP primary antibody for 1.5 h at room temperature (RT) with gentle shaking and consequently with HRP-conjugated secondary anti-mouse antibody (1:5000) for 1 h at RT with gentle shaking. Plaques were visualized using True Blue peroxidase substrate (Sera Care) and the titers were calculated as plaque-forming units per ml (PFU/ml) for all the samples: 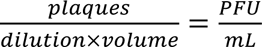

### Intracellular staining for infectivity (Flow cytometry)

ALI cultures were washed with 1X PBS and treated first with 0.05% trypsin on apical and basal chamber (15 min, 37°C). Then, the cells were resuspended in a dispase I/DNase I solution (final concentration in 1X PBS, 1.9U/ml diaspase I (Sigma) and 0,1mg/ml DNase I (Sigma)) to generate single-cell suspensions. The cells were finally resuspended in 1X PBS and stained with zombie live dead stains (Zombie aqua or zombie green; Biolegend) in 1:350 dilution for 15 min at RT. The cells were washed with 1 ml of 1X PBS and finally resuspended in 100μl of 4% methanol-free formaldehyde. The cells were moved out of ABSL3 and left at 4°C for overnight fixation. 1ml of 1X perm wash buffer (BD Biosciences) was added to each tube containing fixed cells and centrifuged at 300g for 5 min. The pellet was resuspended in 100μl perm was buffer with 1:100 dilution of Alexa fluor 488-conjugated or 1:200 of Alexa fluor 647-conjugated SARS-CoV-2 N antibody (IC7) and incubated in dark at RT for 45 min. The cells were washed with 1ml perm wash buffer and centrifuged at 300g for 5 min. The final cell pellet was resuspended in 200μl FACS buffer (1X PBS+ 1% BSA) and analyzed with Flow cytometer for live SARS-CoV-2 NP positive cells.

### Immunofluorescence staining

ALI cultures were fixed using 4% PFA (15 min) *in situ* on inserts, washed with 1X PBS, then embedded in OCT (Sakura Finetek U.S.A.) and snap frozen in liquid nitrogen. Frozen sections were cut at 8 µm, air dried on Superfrost plus slides, fixed again with 4% PFA and permeabilized with 1X PBS/0.1% Triton-X (15 min). Tissue sections were treated with Fc Receptor Block, followed by Background Buster (Innovex Bioscience). The sections were then stained with appropriate primary antibodies (Abs) for one hour followed by appropriate secondary antibodies for 30 min at room temperature in 1X PBS/5% BSA/0.05% saponin. If staining panel included staining with directly conjugated antibodies, tissues were washed, and secondary antibodies were saturated using mouse normal serum diluted at 1/20 in 1X PBS for 15 minutes at room temperature. Then, sections were stained with directly conjugated antibody mix for 1 hour at room temperature and washed. Respective isotype antibodies were used as controls. Finally, sections were counterstained with 1 µg/ml of 4’,6-diamidino-2-phenylindole (DAPI) and 10^-2^ nmol units/ml of Phalloidin ATTO647N (Sigma 65906), mounted with Fluoromount-G (Thermo Fisher Scientific), acquired using a Leica SP8 confocal microscope (for high resolution images) or Thunder widefield microscope (for histocytometry) both with Leica LAS X software and analyzed using Imaris software (Bitplane).

## Quantification and statistical analysis

All results are expressed as mean, unless specified. Descriptive statistics and statistical analysis involved use of Prism 8 (GraphPad Software, San Diego, CA), significance was analyzed by two-tailed non-parametric and unpaired Wilcoxon-Mann-Whitney test using. Statistical significance was determined as P value p ≤ 0.05 (*), p ≤ 0.01(**), p ≤ 0.001 (***) and p ≤ 0.0001 (****). All comparisons were made between infection conditions with time point-matched, uninfected controls.

### GeoMx Whole Transcriptome Atlas slide preparation

#### 1. ALI tissue preparation

ALI cultures were fixed using 4% PFA (15 min) in situ on inserts, washed with 1X PBS, then embedded in OCT (Sakura Finetek U.S.A.) and snap frozen in liquid nitrogen. Frozen sections were cut at 6 µm, air dried on Superfrost plus slides, shipped in dry ice and stored in a secured −80 °C freezer until use. For the NanoString GeoMx^®^ whole transcriptome atlas (WTA) assay that contains 18,676 genes, PFA4% fixed-frozen tissue slides were prepared according to the GeoMx NGS automated Leica Bond RNA Slide Preparation Manual (NanoString, MAN-10131-03).

#### 2. GeoMx DSP instrument and ROI selection

Slides were loaded into the slide holder of the GeoMx digital spatial profiling (DSP) instrument and covered with 2 mL of buffer S. Each slide was scanned with the following antibodies and exposure times: respectively for the morphology markers nuclei Syto83 (Cy3/568nm, 60ms), CK5 (FITC/525nm, 200ms), SARS-CoV-2 Spike (Texas Red/615nm, 250ms), and SARS-CoV-2 Nucleocapsid (Cy5/666nm, 200ms). Corresponding regions of interest (ROIs) were then selected on the DSP slide by their morphology using cytokeratin as basal epithelial cells marker and Spike/Nucleocapside as “infection” marker. In ALI culture selected ROIs were: in mock-infected (control without virus without virus harvested at 6 DPI) tissue, Mock CK5^+^ (CK5^+^ in mock-infected control tissue) and their corresponding Mock CK5^-^ area (Apical area in mock-infected control tissue); in infected tissue identified CK5^+^ staining (CK5^+^ in infected tissue negative for viral markers) and their corresponding CK5^-^ area (Apical area in infected tissue positive for Spike/nucleocapsid viral markers).

After ROIs were approved, the GeoMx DSP exposes the selected regions to UV light which photocleaves the UV cleavable barcode linked from the bound RNA probes, which are collected and deposed into separate wells in the DSP collection plate.

#### 3. GeoMx RNA Illumina library preparation

DSP collection sample plates were dried, resuspended in nuclease-free water, and amplified using PCR according to the manufacturer’s protocol. Purified libraries were sequenced by Illumina NovaSeq6000.

#### 4. GeoMx data processing and QC

The FASTQ reads from sequenced DSP library were processed by the GeoMx NGS DnD Pipeline to convert sequencing reads into counts per ROI (NanoString, MAN-10133_03). After processing, counts were uploaded to the GeoMx DSP Data Analysis Suite (NanoString). QC steps were carried out to assess raw read threshold, percent aligned reads and sequencing saturation. Data was normalized to the third quartile (Q3) to account for differences in cellularity, ROI Size, etc. Analyzing 78 ROIs, all samples had greater than 50% sequencing saturation, 17,265 genes normalized by 3rd quartile were expressed above LOQ in at least 95% of ROIs.

#### 5. GeoMx data analysis

Data analysis was performed in R (v4.2.0). Graphics were produced using ggplot2 unless otherwise stated (v3.3.6).

<u>Differentially expressed genes</u>: A Wilcoxon rank-sum test was used to identify differentially expressed genes between different regions of interest. p values were adjusted using Benjamini-Hochberg multiple test correction. An adjusted p value cut-off of 0.05 and a logFC of 2 was used to identify differentially expressed genes. Variable genes across the whole dataset were identified by a Kruskal-Wallis test (p < 0.05) across the dataset in order to show clustering of different ROI groups by heatmap.

<u>Gene expression heatmaps</u>: Heatmaps were produced using the ComplexHeatmappackage (v2.12.1). Heatmaps used by-row scaling, and ROIs were first grouped by infection type, ordered by Day, and then clustered using the default hierarchical clustering algorithm.

### RNA sequencing and bioinformatics

#### 1. Samples preparation for Bulk RNA sequencing

Total RNA was isolated using Direct-zol RNA MicroPrep kits (zymo research) following the manufacturer’s protocols. During RNA isolation, DNase treatment was additionally performed using the RNase-free DNase set (Qiagen). cDNA libraries were prepared with polyA mRNA selection and Kapa Stranded mRNA-Seq Library Prep kit (Kapa Biosystems) according to the manufacturer’s instructions. Paired-end sequencing (2 × 150 bp) of stranded total RNA libraries was performed in Illumina Novaseq.

#### 2. Bulk RNA sequencing analysis

Illumina was used for demultiplexing. Quality control was performed using FastQC ^93^ (quality control tool for high throughput sequence data) and MultiQC^94^ tools rRNA contamination was removed, and all reads were trimmed using BBDuk tool^95^. Viral reads were mapped to the FDA-ARGOS SARS-CoV-2 reference sequences (FDAARGOS_983, MT233526.1, MT246667.1; EPI_ISL_10489564 was obtained from GISAID database^96^), and human reads were mapped to GRCh38 genome using bowtie^97,98^. Gene expression was quantified using RSEM^99^. Batch correction was removed using R package SVAseq^100^. Reads were normalized and differentially expressed genes were analyzed using DESeq2^101^. Pathway analysis was performed using QIAGEN IPA^102,103^

## Supplemental information

### Document with supplementary figures S1-S6

**Table S1 (Related to Figure 1): Demographic information of lung donors:** Epithelial lung organoids from cryopreserved lung pieces from four healthy lung donors (all Hispanic, 3 male donors aged between 34-50 y/o and 1 female donor 54 y/o).

**Table S2:**

- **Sheet 1 (Related to figure 1): Descriptive statistics of histocytometry data of cell markers expression by primary ALI cultures.** Four major epithelial markers characterized ALI cultures: “Acetyl” for acetyl-alpha-tubulin marker related to ciliated cells; “SCGB1A1” marker related to club cells; “Muc5ac” marker related to goblet cells and “CK5” for cytokeratin 5 related to basal cells. The “Other” category corresponded to the cells that did not express any of the given markers. Data shown are representatives of 3 independent experiments per donor with 1 replicate per experiment. Normalized histograms and descriptive statistics were generated using Prism 8 Graph Pad.
- **Sheet 2 (Related to figure 2): Descriptive statistics of histocytometry, flow cytometry and viral titer data of primary ALI cultures exposed to SARS-CoV-2 Wuhan-like (USA/WA1-2020).** Data shown are representatives of 9-10 independent experiments, at least 2 independent experiments per donor with 1 replicate per experiment. Descriptive statistics were generated using Prism 8 Graph Pad.
- **Sheet 3 (Related to figure 3): Descriptive statistics of histocytometry data of primary ALI cultures exposed to SARS-CoV-2 variants.** Data shown are representatives of at least 2-3 independent experiments per variant, 1 replicate per experiment. Descriptive statistics were generated using Prism 8 Graph Pad.
- **Sheet 4 (Related to Figure S3): Descriptive statistics of flow cytometry and viral titer data of primary ALI cultures exposed to SARS-CoV-2 variants.** Data shown are representatives of at least 2-3 independent experiments per variant. Descriptive statistics were generated using Prism 8 Graph Pad.

**Table S3 (Related to Figure 4):** differentially expressed genes (DEG) list values for SARS-CoV-2 Wuhan-like virus data.

**Table S4 (Related to Figure 5):** differentially expressed genes (DEG) list values for SARS-CoV-2 variants data, statistics, and descriptive statistics for line plots were generated using Prism 8 Graph Pad.

**Table S5 (Related to Figure 7):** GeoMx DSP analysis, number of regions of interest (ROI) selected per tissue.

**Table S6 (Related to Figure 7):** GeoMx gene list analysis for SARS-CoV-2 Wuhan-like virus.

## References

1. Blanco-Melo, D., Nilsson-Payant, B.E., Liu, W.C., Uhl, S., Hoagland, D., Møller, R., Jordan, T.X., Oishi, K., Panis, M., Sachs, D., et al. (2020). Imbalanced Host Response to SARS-CoV-2 Drives Development of COVID-19. Cell 181, 1036–1045.e9. 10.1016/j.cell.2020.04.026.

2. Flerlage, T., Boyd, D.F., Meliopoulos, V., Thomas, P.G., and Schultz-Cherry, S. (2021). Influenza virus and SARS-CoV-2: pathogenesis and host responses in the respiratory tract. Nat. Rev. Microbiol. 19, 425–441. 10.1038/s41579-021-00542-7.

3. Hartshorn, K.L. (2020). Innate Immunity and Influenza A Virus Pathogenesis: Lessons for COVID-19. Front. Cell. Infect. Microbiol. 10, 1–17. 10.3389/fcimb.2020.563850.

4. Mangalmurti, N., and Hunter, C.A. (2020). Cytokine Storms: Understanding COVID-19. Immunity 53, 19–25. 10.1016/j.immuni.2020.06.017.

5. Niec, R.E., Rudensky, A.Y., and Fuchs, E. (2021). Inflammatory adaptation in barrier tissues. Cell 184, 3361–3375. 10.1016/j.cell.2021.05.036.

6. Ziegler, C.G.K., Allon, S.J., Nyquist, S.K., Mbano, I.M., Miao, V.N., Tzouanas, C.N., Cao, Y., Yousif, A.S., Bals, J., Hauser, B.M., et al. (2020). SARS-CoV-2 Receptor ACE2 Is an Interferon-Stimulated Gene in Human Airway Epithelial Cells and Is Detected in Specific Cell Subsets across Tissues. Cell 181, 1016–1035.e19. 10.1016/j.cell.2020.04.035.

7. Klouda, T., Hao, Y., Kim, H., Kim, J., Olejnik, J., Hume, A.J., Ayyappan, S., Hong, X., Melero-Martin, J., Fang, Y., et al. (2021). Interferon-alpha or -beta facilitates SARS-CoV-2 pulmonary vascular infection by inducing ACE2. Angiogenesis. 10.1007/s10456-021-09823-4.

8. Zhou, Y., Wang, M., Li, Y., Wang, P., Zhao, P., Yang, Z., Wang, S., Zhang, L., Li, Z., Jia, K., et al. (2021). SARS-CoV-2 Spike protein enhances ACE2 expression via facilitating Interferon effects in bronchial epithelium. Immunol. Lett. 237, 33–41. 10.1016/j.imlet.2021.06.008.

9. V’kovski, P., Kratzel, A., Steiner, S., Stalder, H., and Thiel, V. (2021). Coronavirus biology and replication: implications for SARS-CoV-2. Nat. Rev. Microbiol. 19, 155–170. 10.1038/s41579-020-00468-6.

10. Lukassen, S., Chua, R.L., Trefzer, T., Kahn, N.C., Schneider, M.A., Muley, T., Winter, H., Meister, M., Veith, C., Boots, A.W., et al. (2020). SARS-CoV-2 receptor ACE 2 and TMPRSS 2 are primarily expressed in bronchial transient secretory cells. EMBO J. 39. 10.15252/embj.20105114.

11. Hoffmann, M., Kleine-Weber, H., Schroeder, S., Krüger, N., Herrler, T., Erichsen, S., Schiergens, T.S., Herrler, G., Wu, N.H., Nitsche, A., et al. (2020). SARS-CoV-2 Cell Entry Depends on ACE2 and TMPRSS2 and Is Blocked by a Clinically Proven Protease Inhibitor. Cell 181, 271–280.e8. 10.1016/j.cell.2020.02.052.

12. Peng, R., Wu, L.A., Wang, Q., Qi, J., and Gao, G.F. (2021). Cell entry by SARS-CoV-2. Trends Biochem. Sci. 46, 848–860. 10.1016/j.tibs.2021.06.001.

13. Hasan, M.Z., Islam, S., Matsumoto, K., and Kawai, T. (2021). SARS-CoV-2 infection initiates interleukin-17-enriched transcriptional response in different cells from multiple organs. Sci. Rep. 11, 1–11. 10.1038/s41598-021-96110-3.

14. Chua, R.L., Lukassen, S., Trump, S., Hennig, B.P., Wendisch, D., Pott, F., Debnath, O., Thürmann, L., Kurth, F., Völker, M.T., et al. (2020). COVID-19 severity correlates with airway epithelium–immune cell interactions identified by single-cell analysis. Nat. Biotechnol. 38, 970–979. 10.1038/s41587-020-0602-4.

15. Ackermann, M., Anders, H.J., Bilyy, R., Bowlin, G.L., Daniel, C., De Lorenzo, R., Egeblad, M., Henneck, T., Hidalgo, A., Hoffmann, M., et al. (2021). Patients with COVID-19: in the dark-NETs of neutrophils. Cell Death Differ. 28, 3125–3139. 10.1038/S41418-021-00805-Z.

16. Channappanavar, R., Fehr, A.R., Vijay, R., Mack, M., Zhao, J., Meyerholz, D.K., and Perlman, S. (2016). Dysregulated Type I Interferon and Inflammatory Monocyte-Macrophage Responses Cause Lethal Pneumonia in SARS-CoV-Infected Mice. Cell Host Microbe 19, 181–193. 10.1016/j.chom.2016.01.007.

17. Boumaza, A., Gay, L., Mezouar, S., Bestion, E., Diallo, A.B., Michel, M., Desnues, B., Raoult, D., La Scola, B., Halfon, P., et al. (2021). Monocytes and macrophages, targets of SARS-CoV-2: the clue for Covid-19 immunoparalysis. J. Infect. Dis. 224, 395–406. 10.1093/INFDIS/JIAB044.

18. Leon, J., Michelson, D.A., Olejnik, J., Chowdhary, K., Oh, H.S., Hume, A.J., Galvan-Peña, S., Zhu, Y., Chen, F., Vijaykumar, B., et al. (2022). A virus-specific monocyte inflammatory phenotype is induced by SARS-CoV-2 at the immune– epithelial interface. Proc. Natl. Acad. Sci. U. S. A. 119. 10.1073/pnas.2116853118.

19. Ribero, M.S., Jouvenet, N., Dreux, M., and Nisole, S. (2020). Interplay between SARS-CoV-2 and the type I interferon response. PLoS Pathog. 16. 10.1371/JOURNAL.PPAT.1008737.

20. Laurent, P., Yang, C., Rendeiro, A.F., Nilsson-Payant, B.E., Carrau, L., Chandar, V., Bram, Y., tenOever, B.R., Elemento, O., Ivashkiv, L.B., et al. (2022). Sensing of SARS-CoV-2 by pDCs and their subsequent production of IFN-I contribute to macrophage-induced cytokine storm during COVID-19. Sci. Immunol. 7, eadd4906. 10.1126/SCIIMMUNOL.ADD4906/SUPPL_FILE/SCIIMMUNOL.ADD4906_TABLES_S1_TO_S6.ZIP.

21. Afzali, B., Noris, M., Lambrecht, B.N., and Kemper, C. (2021). The state of complement in COVID-19. Nat. Rev. Immunol. 2021 222 22, 77–84. 10.1038/s41577-021-00665-1.

22. Ma, L. (2021). Increased complement activation is a distinctive feature of severe SARS-CoV-2 infection. Sci. Immunol. 6.

23. Ramlall, V. (2020). Immune complement and coagulation dysfunction in adverse outcomes of SARS-CoV-2 infection. Nat. Med. 26, 1609–1615.

24. Sinkovits, G., Schnur, J., Hurler, L., Kiszel, P., Prohászka, Z.Z., Sík, P., Kajdácsi, E., Cervenak, L., Maráczi, V., Dávid, M., et al. (2022). Evidence, detailed characterization and clinical context of complement activation in acute multisystem inflammatory syndrome in children. Sci. Rep. 12, 19759. 10.1038/S41598-022-23806-5.

25. Shah, V.K., Firmal, P., Alam, A., Ganguly, D., and Chattopadhyay, S. (2020). Overview of Immune Response During SARS-CoV-2 Infection: Lessons From the Past. Front. Immunol. 11, 1949. 10.3389/FIMMU.2020.01949/XML/NLM.

26. Cox, R.J., and Brokstad, K.A. (2020). Not just antibodies: B cells and T cells mediate immunity to COVID-19. Nat. Rev. Immunol. 2020 2010 20, 581–582. 10.1038/s41577-020-00436-4.

27. Sterlin, D., Mathian, A., Miyara, M., Mohr, A., Anna, F., Claër, L., Quentric, P., Fadlallah, J., Devilliers, H., Ghillani, P., et al. (2021). IgA dominates the early neutralizing antibody response to SARS-CoV-2. Sci. Transl. Med. 13, 2223. 10.1126/SCITRANSLMED.ABD2223.

28. Khan, A., Khan, S.A., Zia, K., Altowyan, M.S., Barakat, A., and Ul-Haq, Z. (2022). Deciphering the Impact of Mutations on the Binding Efficacy of SARS-CoV-2 Omicron and Delta Variants With Human ACE2 Receptor. Front. Chem. 10, 566. 10.3389/FCHEM.2022.892093/BIBTEX.

29. Khateeb, J., Li, Y., and Zhang, H. (2021). Emerging SARS-CoV-2 variants of concern and potential intervention approaches. Crit. Care 25, 1–8. 10.1186/S13054-021-03662-X/TABLES/2.

30. Kumar, S., Thambiraja, T.S., Karuppanan, K., and Subramaniam, G. (2022). Omicron and Delta variant of SARS-CoV-2: A comparative computational study of spike protein. J. Med. Virol. 94, 1641–1649. 10.1002/jmv.27526.

31. Reincke, S.M., Yuan, M., Kornau, H.C., Corman, V.M., van Hoof, S., Sánchez-Sendin, E., Ramberger, M., Yu, W., Hua, Y., Tien, H., et al. (2022). SARS-CoV-2 Beta variant infection elicits potent lineage-specific and cross-reactive antibodies. Science (80-.). 375, 782–787. 10.1126/SCIENCE.ABM5835/SUPPL_FILE/SCIENCE.ABM5835_SM_MDAR_REPRODUCIBILITY_CHECKLIST.PDF.

32. Rajah, M.M., Hubert, M., Bishop, E., Saunders, N., Robinot, R., Grzelak, L., Planas, D., Dufloo, J., Gellenoncourt, S., Bongers, A., et al. (2021). SARS-CoV-2 Alpha, Beta, and Delta variants display enhanced Spike-mediated syncytia formation. EMBO J. 40. 10.15252/EMBJ.2021108944.

33. Suzuki, R., Yamasoba, D., Kimura, I., Wang, L., Kishimoto, M., Ito, J., Morioka, Y., Nao, N., Nasser, H., Uriu, K., et al. (2022). Attenuated fusogenicity and pathogenicity of SARS-CoV-2 Omicron variant. 603, 700–705.

34. Lamers, M.M., Mykytyn, A.Z., Breugem, T.I., Groen, N., Knoops, K., Schipper, D., Acker, R. van, Doel, P.B. van den, Bestebroer, T., Koopman, C.D., et al. (2022). SARS-CoV-2 Omicron efficiently infects human airway, but not alveolar epithelium. bioRxiv, 2022.01.19.476898.

35. Hui, K.P.Y., Ho, J.C.W., Cheung, M. chun, Ng, K. chun, Ching, R.H.H., Lai, K. ling, Kam, T.T., Gu, H., Sit, K.Y., Hsin, M.K.Y., et al. (2022). SARS-CoV-2 Omicron variant replication in human bronchus and lung ex vivo. Nat. 2022 6037902 603, 715–720. 10.1038/s41586-022-04479-6.

36. Pia, L., and Rowland-Jones, S. (2022). Omicron entry route. Nat. Rev. Immunol. 2022 223 22, 144–144. 10.1038/s41577-022-00681-9.

37. Shrestha, L.B., Foster, C., Rawlinson, W., Tedla, N., and Bull, R.A. (2022). Evolution of the SARS-CoV-2 omicron variants BA.1 to BA.5: Implications for immune escape and transmission. Rev. Med. Virol. 32, 32. 10.1002/RMV.2381.

38. Aggarwal, A., Akerman, A., Milogiannakis, V., Silva, M.R., Walker, G., Stella, A.O., Kindinger, A., Angelovich, T., Waring, E., Amatayakul-Chantler, S., et al. (2022). SARS-CoV-2 Omicron BA.5: Evolving tropism and evasion of potent humoral responses and resistance to clinical immunotherapeutics relative to viral variants of concern. eBioMedicine 84, 104270. 10.1016/j.ebiom.2022.104270.

39. Fulcher, M.L., Gabriel, S., Burns, K.A., Yankaskas, J.R., and Randell, S.H. (2005). Well-differentiated human airway epithelial cell cultures. Methods Mol. Med. 10.1385/1-59259-861-7:183.

40. Rayner, R.E., Makena, P., Prasad, G.L., and Cormet-Boyaka, E. (2019). Optimization of Normal Human Bronchial Epithelial (NHBE) Cell 3D Cultures for in vitro Lung Model Studies. Sci. Rep. 10.1038/s41598-018-36735-z.

41. Sachs, N., Papaspyropoulos, A., Zomer-van Ommen, D.D., Heo, I., Böttinger, L., Klay, D., Weeber, F., Huelsz-Prince, G., Iakobachvili, N., Amatngalim, G.D., et al. (2019). Long-term expanding human airway organoids for disease modeling. EMBO J. 38, 1–20. 10.15252/embj.2018100300.

42. Srinivasan, B., Kolli, A.R., Esch, M.B., Abaci, H.E., Shuler, M.L., and Hickman, J.J. (2015). TEER Measurement Techniques for In Vitro Barrier Model Systems. J. Lab. Autom. 10.1177/2211068214561025.

43. Brest, P., Refae, S., Mograbi, B., Hofman, P., and Milano, G. (2020). Host Polymorphisms May Impact SARS-CoV-2 Infectivity. Trends Genet. 36, 813–815. 10.1016/j.tig.2020.08.003.

44. Adli, A., Rahimi, M., Khodaie, R., Hashemzaei, N., and Hosseini, S.M. (2022). Role of genetic variants and host polymorphisms on COVID-19: From viral entrance mechanisms to immunological reactions. J. Med. Virol. 94, 1846. 10.1002/JMV.27615.

45. Peng, G., Ke, J.L., Jin, W., Greenwell-Wild, T., and Wahl, S.M. (2006). Induction of APOBEC3 family proteins, a defensive maneuver underlying interferon-induced anti-HIV-1 activity. J. Exp. Med. 203, 41–46. 10.1084/JEM.20051512.

46. Milewska, A., Kindler, E., Vkovski, P., Zeglen, S., Ochman, M., Thiel, V., Rajfur, Z., and Pyrc, K. (2018). APOBEC3-mediated restriction of RNA virus replication. Sci. Rep. 8. 10.1038/S41598-018-24448-2.

47. Coperchini, F., Chiovato, L., Ricci, G., Croce, L., Magri, F., and Rotondi, M. (2021). The cytokine storm in COVID-19: Further advances in our understanding the role of specific chemokines involved. Cytokine Growth Factor Rev. 58, 82–91. 10.1016/j.cytogfr.2020.12.005.

48. Nunnari, G., Sanfilippo, C., Castrogiovanni, P., Imbesi, R., Li Volti, G., Barbagallo, I., Musumeci, G., and Di Rosa, M. (2020). Network perturbation analysis in human bronchial epithelial cells following SARS-CoV2 infection. Exp. Cell Res. 395, 112204. 10.1016/J.YEXCR.2020.112204.

49. Hernández-Santos, N., Wiesner, D.L., Fites, J.S., McDermott, A.J., Warner, T., Wüthrich, M., and Klein, B.S. (2018). Lung Epithelial Cells Coordinate Innate Lymphocytes and Immunity against Pulmonary Fungal Infection. Cell Host Microbe 23, 511–522.e5. 10.1016/j.chom.2018.02.011.

50. Smieszek, S.P., Polymeropoulos, V.M., Polymeropoulos, C.M., Przychodzen, B.P., Birznieks, G., and Polymeropoulos, M.H. (2022). Elevated plasma levels of CXCL16 in severe COVID-19 patients. Cytokine 152, 155810. 10.1016/j.cyto.2022.155810.

51. Fricke-Galindo, I., and Falfán-Valencia, R. (2021). Genetics Insight for COVID-19 Susceptibility and Severity: A Review. Front. Immunol. 12, 1–11. 10.3389/fimmu.2021.622176.

52. Rovito, R., Augello, M., Ben-Haim, A., Bono, V., d’Arminio Monforte, A., and Marchetti, G. (2022). Hallmarks of Severe COVID-19 Pathogenesis: A Pas de Deux Between Viral and Host Factors. Front. Immunol. 13, 2576. 10.3389/FIMMU.2022.912336/BIBTEX.

53. Hsu, R.J., Yu, W.C., Peng, G.R., Ye, C.H., Hu, S.Y., Chong, P.C.T., Yap, K.Y., Lee, J.Y.C., Lin, W.C., and Yu, S.H. (2022). The Role of Cytokines and Chemokines in Severe Acute Respiratory Syndrome Coronavirus 2 Infections. Front. Immunol. 13. 10.3389/FIMMU.2022.832394/FULL.

54. Vanderheiden, A., Ralfs, P., Chirkova, T., Upadhyay, A.A., Zimmerman, M.G., Bedoya, S., Aoued, H., Tharp, G.M., Pellegrini, K.L., Manfredi, C., et al. (2020). Type I and Type III Interferons Restrict SARS-CoV-2 Infection of Human Airway Epithelial Cultures. J. Virol. 94, 1–16. 10.1128/jvi.00985-20.

55. Hadjadj, J., Yatim, N., Barnabei, L., Corneau, A., and Boussier, J. (2020). Impaired type I IFN activity and inflammation. 724, 718–724.

56. Roth, W., Kumar, V., Beer, H.-D., Richter, M., Wohlenberg, C., Reuter, U., Ren Thiering, S., Staratschek-Jox, A., Hofmann, A., Kreusch, F., et al. Keratin 1 maintains skin integrity and participates in an inflammatory network in skin through interleukin-18. J. Cell Sci. 125, 5269–5279. 10.1242/jcs.116574.

57. Boldt, K., Van Reeuwijk, J., Lu, Q., Koutroumpas, K., Nguyen, T.M.T., Texier, Y., Van Beersum, S.E.C., Horn, N., Willer, J.R., Mans, D.A., et al. (2016). An organelle-specific protein landscape identifies novel diseases and molecular mechanisms. Nat. Commun. 2016 71 7, 1–13. 10.1038/ncomms11491.

58. Chagula, D.B., Rechciński, T., Rudnicka, K., and Chmiela, M. (2020). Ankyrins in human health and disease – an update of recent experimental findings. Arch. Med. Sci. 16, 715. 10.5114/AOMS.2019.89836.

59. Sánchez-Navarro, A., González-Soria, I., Caldi∼ No-Bohn, R., and Bobadilla, N.A. (2021). An integrative view of serpins in health and disease: the contribution of SerpinA3. 10.1152/ajpcell.00366.2020.

60. Ma, X., Hua, J., Zheng, G., Li, F., Rao, C., Li, H., Wang, J., Pan, L., and Hou, L. (2018). Regulation of cell proliferation in the retinal pigment epithelium: Differential regulation of the death-associated protein like-1 DAPL1 by alternative MITF splice forms. Pigment Cell Melanoma Res. 31, 411–422. 10.1111/PCMR.12676.

61. Ross, A.J., Dailey, L.A., Brighton, L.E., and Devlin, R.B. (2012). Transcriptional Profiling of Mucociliary Differentiation in Human Airway Epithelial Cells. https://doi.org/10.1165/rcmb.2006-0466OC 37, 169–185. 10.1165/RCMB.2006-0466OC.

62. Nanostring (2017). Explore the biology that matters Spatial RNA profiling aligns with RNASeq and RNAscope.

63. GeoMx DSP Spatial Genomics Overview - NanoString https://nanostring.com/products/geomx-digital-spatial-profiler/geomx-dsp-overview/?utm_source=AdWords&utm_medium=SearchAds&utm_campaign=DSP&gclid=CjwKCAiApvebBhAvEiwAe7mHSHqddg31w0IOdaJm5WYksuOsvbN04omovyvZTqEbe5_wf5PihOLdkxoCwzsQAvD_BwE.

64. Sheng, H., Shao, J., Townsend, C.M., and Evers, B.M. (2003). Phosphatidylinositol 3-kinase mediates proliferative signals in intestinal epithelial cells. Gut 52, 1472. 10.1136/GUT.52.10.1472.

65. Barratt, S.L., Blythe, T., Jarrett, C., Ourradi, K., Shelley-Fraser, G., Day, M.J., Qiu, Y., Harper, S., Maher, T.M., Oltean, S., et al. (2017). Differential Expression of VEGF-A xxx Isoforms Is Critical for Development of Pulmonary Fibrosis. 10.1164/rccm.201603-0568OC.

66. Besnard, V., Corroyer, S., Trugnan, G., Chadelat, K., Nabeyrat, E., Cazals, V., and Clement, A. (2001). Distinct patterns of insulin-like growth factor binding protein (IGFBP)-2 and IGFBP-3 expression in oxidant exposed lung epithelial cells. Biochim. Biophys. Acta 1538, 47–58. 10.1016/S0167-4889(00)00136-1.

67. Yoshioka, M., Sawada, Y., Saito-Sasaki, N., Yoshioka, H., Hama, K., Omoto, D., Ohmori, S., Okada, E., and Nakamura, M. (2021). High S100A2 expression in keratinocytes in patients with drug eruption. Sci. Reports 2021 111 11, 1–9. 10.1038/s41598-021-85009-8.

68. Ghosh, M., Ahmad, S., Jian, A., Li, B., Smith, R.W., Helm, K.M., Seibold, M.A., Groshong, S.D., White, C.W., and Reynolds, S.D. (2013). Human tracheobronchial basal cells: Normal versus remodeling/repairing phenotypes in vivo and in vitro. Am. J. Respir. Cell Mol. Biol. 49, 1127–1134. 10.1165/RCMB.2013-0049OC/SUPPL_FILE/DISCLOSURES.PDF.

69. Hewitt, R.J., and Lloyd, C.M. (2021). Regulation of immune responses by the airway epithelial cell landscape. Nat. Rev. Immunol. 21, 347–362. 10.1038/s41577-020-00477-9.

70. Garcıá, S.R., Deprez, M., Lebrigand, K., Cavard, A., Paquet, A., Arguel, M.J., Magnone, V., Truchi, M., Caballero, I., Leroy, S., et al. (2019). Novel dynamics of human mucociliary differentiation revealed by single-cell RNA sequencing of nasal epithelial cultures. Development 146. 10.1242/DEV.177428.

71. Gasser, M., and Waaga-Gasser, A.M. (2016). Therapeutic Antibodies in Cancer Therapy. In (Springer, Cham), pp. 95–120.

72. Smirnova, N.F., Schamberger, A.C., Nayakanti, S., Hatz, R., Behr, J., and Eickelberg, O. (2016). Detection and quantification of epithelial progenitor cell populations in human healthy and IPF lungs. Respir. Res. 17, 1–11. 10.1186/S12931-016-0404-X/FIGURES/8.

73. Tran, B.M., Deliyannis, G., Hachani, A., Earnest, L., Torresi, J., and Vincan, E. (2022). Organoid Models of SARS-CoV-2 Infection: What Have We Learned about COVID-19? Organoids 2022, Vol. 1, Pages 2-27 1, 2–27. 10.3390/ORGANOIDS1010002.

74. Willett, B.J., Grove, J., MacLean, O.A., Wilkie, C., De Lorenzo, G., Furnon, W., Cantoni, D., Scott, S., Logan, N., Ashraf, S., et al. (2022). SARS-CoV-2 Omicron is an immune escape variant with an altered cell entry pathway. Nat. Microbiol. 2022 78 7, 1161–1179. 10.1038/s41564-022-01143-7.

75. Peacock, T.P., Brown, J.C., Zhou, J., Thakur, N., Newman, J., Kugathasan, R., Sukhova, K., Kaforou, M., Bailey, D., and Barclay, W.S. (2022). The SARS-CoV-2 variant, Omicron, shows rapid replication in human primary nasal epithelial cultures and efficiently uses the endosomal route of entry. bioRxiv, 2021.12.31.474653. 10.1101/2021.12.31.474653.

76. Hao, S., Ning, K., Kuz, C.A., Vorhies, K., Yan, Z., and Qiu, J. (2020). Long-Term Modeling of SARS-CoV-2 Infection of. Am. J. Microbiol. 11, 1–17.

77. Kikutani, H., and Kumanogoh, A. (2003). Semaphorins in interactions between T cells and antigen-presenting cells. Nat. Rev. Immunol. 2003 32 3, 159–167. 10.1038/nri1003.

78. Wang, H., FitzPatrick, M., Wilson, N.J., Anthony, D., Reading, P.C., Satzke, C., Dunne, E.M., Licciardi, P. V., Seow, H.J., Nichol, K., et al. (2019). CSF3R/CD114 mediates infection-dependent transition to severe asthma. J. Allergy Clin. Immunol. 143, 785–788.e6. 10.1016/j.jaci.2018.10.001.

79. Touzelet, O., Broadbent, L., Armstrong, S.D., Aljabr, W., Cloutman-Green, E., Power, U.F., and Hiscox, J.A. (2020). The Secretome Profiling of a Pediatric Airway Epithelium Infected with hRSV Identified Aberrant Apical/Basolateral Trafficking and Novel Immune Modulating (CXCL6, CXCL16, CSF3) and Antiviral (CEACAM1) Proteins. Mol. Cell. Proteomics 19, 793. 10.1074/MCP.RA119.001546.

80. Fang, C., Mei, J., Tian, H., Liou, Y.L., Rong, D., Zhang, W., Liao, Q., and Wu, N. (2020). CSF3 Is a Potential Drug Target for the Treatment of COVID-19. Front. Physiol. 11. 10.3389/FPHYS.2020.605792.

81. Klein, M., Brouwer, M.C., Angele, B., Geldhoff, M., Marquez, G., Varona, R., Häcker, G., Schmetzer, H., Häcker, H., Hammerschmidt, S., et al. (2014). Leukocyte Attraction by CCL20 and Its Receptor CCR6 in Humans and Mice with Pneumococcal Meningitis. PLoS One 9, e93057. 10.1371/JOURNAL.PONE.0093057.

82. Robinot, R., Hubert, M., de Melo, G.D., Lazarini, F., Bruel, T., Smith, N., Levallois, S., Larrous, F., Fernandes, J., Gellenoncourt, S., et al. (2021). SARS-CoV-2 infection induces the dedifferentiation of multiciliated cells and impairs mucociliary clearance. Nat. Commun. 2021 121 12, 1–16. 10.1038/s41467-021-24521-x.

83. Schreiner, T., Allnoch, L., Beythien, G., Marek, K., Becker, K., Schaudien, D., Stanelle-Bertram, S., Schaumburg, B., Kouassi, N.M., Beck, S., et al. (2022). SARS-CoV-2 Infection Dysregulates Cilia and Basal Cell Homeostasis in the Respiratory Epithelium of Hamsters. Int. J. Mol. Sci. 23, 5124. 10.3390/IJMS23095124/S1.

84. Nilsson-Payant, B.E., Uhl, S., Grimont, A., Doane, A.S., Cohen, P., Patel, R.S., Higgins, C.A., Acklin, J.A., Bram, Y., Chandar, V., et al. (2021). The NF-κB Transcriptional Footprint Is Essential for SARS-CoV-2 Replication. J. Virol. 95. 10.1128/JVI.01257-21.

85. Mache, C., Schulze, J., Holland, G., Bourquain, D., Gensch, J.M., Oh, D.Y., Nitsche, A., Dürrwald, R., Laue, M., and Wolff, T. (2022). SARS-CoV-2 Omicron variant is attenuated for replication in a polarized human lung epithelial cell model. Commun. Biol. 2022 51 5, 1–8. 10.1038/s42003-022-04068-3.

86. Wang, X., Wang, J., Zheng, H., Xie, M., Hopewell, E.L., Albrecht, R.A., Nogusa, S., García-Sastre, A., Balachandran, S., and Beg, A.A. (2014). Differential requirement for the IKKβ/NF-κB signaling module in regulating TLR-versus RLR-induced type 1 IFN expression in dendritic cells. J. Immunol. 193, 2538–2545. 10.4049/JIMMUNOL.1400675.

87. Nies, J.F., and Panzer, U. (2020). IL-17C/IL-17RE: Emergence of a Unique Axis in TH17 Biology. Front. Immunol. 11, 341. 10.3389/FIMMU.2020.00341/BIBTEX.

88. Peng, T., Chanthaphavong, R.S., Sun, S., Trigilio, J.A., Phasouk, K., Jin, L., Layton, E.D., Li, A.Z., Correnti, C.E., De van der Schueren, W., et al. (2017). Keratinocytes produce IL-17c to protect peripheral nervous systems during human HSV-2 reactivation. J. Exp. Med. 214, 2315–2329. 10.1084/JEM.20160581.

89. Bojkova, D., Rothenburger, T., Ciesek, S., Wass, M.N., Michaelis, M., and Cinatl, J. (2022). SARS-CoV-2 Omicron variant virus isolates are highly sensitive to interferon treatment. Cell Discov. 2022 81 8, 1–4. 10.1038/s41421-022-00408-z.

90. Yin, X., Riva, L., Pu, Y., Martin-Sancho, L., Kanamune, J., Yamamoto, Y., Sakai, K., Gotoh, S., Miorin, L., De Jesus, P.D., et al. (2021). MDA5 Governs the Innate Immune Response to SARS-CoV-2 in Lung Epithelial Cells. Cell Rep. 34. 10.1016/J.CELREP.2020.108628.

91. Cao, X., Coyle, J.P., Xiong, R., Wang, Y., Heflich, R.H., Ren, B., Gwinn, W.M., Hayden, P., and Rojanasakul, L. (2021). Invited review: human air-liquid-interface organotypic airway tissue models derived from primary tracheobronchial epithelial cells—overview and perspectives. Vitr. Cell. Dev. Biol. - Anim. 57, 104–132. 10.1007/s11626-020-00517-7.

92. Rathnasinghe, R., Jangra, S., Ye, C., Cupic, A., Singh, G., Martínez-Romero, C., Mulder, L.C.F., Kehrer, T., Yildiz, S., Choi, A., et al. (2022). Characterization of SARS-CoV-2 Spike mutations important for infection of mice and escape from human immune sera. Nat. Commun. 2022 131 13, 1–14. 10.1038/s41467-022-30763-0.

93. Andrews S (2010). FastQC A Quality Control tool for High Throughput Sequence Data. https://www.bioinformatics.babraham.ac.uk/projects/fastqc/.

94. Ewels, P., Magnusson, M., Lundin, S., and Käller, M. (2016). MultiQC: summarize analysis results for multiple tools and samples in a single report. Bioinformatics 32, 3047–3048. 10.1093/BIOINFORMATICS/BTW354.

95. BBMap download | SourceForge.net https://sourceforge.net/projects/bbmap/.

96. Khare, S., Gurry, C., Freitas, L., Schultz, M.B., Bach, G., Diallo, A., Akite, N., Ho, J., Lee, R.T.C., Yeo, W., et al. (2021). GISAID’s Role in Pandemic Response. China CDC Wkly. 3, 1049–1051. 10.46234/CCDCW2021.255.

97. Bowtie: An ultrafast, memory-efficient short read aligner http://bowtie-bio.sourceforge.net/index.shtml.

98. Langmead, B., Trapnell, C., Pop, M., and Salzberg, S.L. (2009). Ultrafast and memory-efficient alignment of short DNA sequences to the human genome. Genome Biol. 10, 1–10. 10.1186/GB-2009-10-3-R25/TABLES/5.

99. Li, B., and Dewey, C.N. (2011). RSEM: Accurate transcript quantification from RNA-Seq data with or without a reference genome. BMC Bioinformatics 12, 1–16. 10.1186/1471-2105-12-323/TABLES/6.

100. Zhang, Y., Parmigiani, G., and Johnson, W.E. (2020). ComBat-seq: batch effect adjustment for RNA-seq count data. NAR Genomics Bioinforma. 2. 10.1093/NARGAB/LQAA078.

101. Love, M.I., Huber, W., and Anders, S. (2014). Moderated estimation of fold change and dispersion for RNA-seq data with DESeq2. Genome Biol. 15, 1–21. 10.1186/S13059-014-0550-8/FIGURES/9.

102. Ingenuity Pathway Analysis | QIAGEN Digital Insights https://digitalinsights.qiagen.com/products-overview/discovery-insights-portfolio/analysis-and-visualization/qiagen-ipa/.

103. Krämer, A., Green, J., Pollard, J., and Tugendreich, S. (2014). Causal analysis approaches in Ingenuity Pathway Analysis. Bioinformatics 30, 523–530. 10.1093/BIOINFORMATICS/BTT703.

