## Supplementary figures for "Spatiotemporally organized immunomodulatory response to SARS-CoV-2 virus in primary human broncho-alveolar epithelia"

### Supplemental information

##### Figure S1 Primary human lung organoids and lung organoid-derived air-liquid interface (ALI) culture. (Related to figure 1)

(A) Representative immunofluorescent (IF) images of whole mounted lung organoids showing markers for epithelial cells (PAN-CK, green), extracellular matrix (Fibronectin, red), and nuclei (DAPI, blue). Pan-CK is present exclusively in epithelial cells and fibronectin allow to visualize the presence of remaining extracellular matrix from tissue dissociation. The fibronectin disappears after four passages and characterize the enrichment of lung epithelial cells. Scale bar 100 μm, in white on the left corner. (B) Representative immunofluorescent (IF) section (8 μm) of differentiated lung organoid-derived ALI cultures of three donors. Left panel, merged figures, showing markers for ciliated cells (acetylated a-tubulin, red) and goblet cells (MUC5AC, cyan). Right panel, merged figures, showing club cells (SCGB1A1, green), basal cells (CK5, white). Nuclei (Dapi, blue). Scale bar 40 μm, in white on the left corner. (C) Measurement of trans-epithelial electrical resistance (TEER, ohm/cm2) with error bars (mean ± SD), 3 measurement per time-point performed 3 times per week starting when the epithelium is confluent, one representative experiment per donor (four donors). A fully pseudostratified differentiated ALI is obtained within 3-4 weeks from primary lung organoid progenitors.

##### Figure S2 SARS-CoV-2 Wuhan-like viral infection assessment of lung organoid-derived air-liquid interface (ALI) cultures by multiple-methods. (Related to figure 2)

(A) Schematic experimental design for primary human lung organoid-derived ALI cultures generation. Primary human lung cell suspension was generated from viable lung freeze tissue (10% DMSO in Fetal Bovine Serum, FBS) and then, organoid and ALI cultures were generated as described in material and methods. Once, the ALI cultures were fully differentiated, SARS-CoV-2 was applied apically ($10^5$ PFU/well) and the cultures were harvested every day after in a kinetic fashion from 1 to 6 DPI. (B) Gating strategy example for flow cytometry analysis. Single cell suspension was prepared from mock-infected (media without virus) and SARS-CoV-2 infected ALI cultures. Cell suspensions were stained for viability and viral infection by using an anti-NP antibody specific to SARS-CoV-2. From left to right, the first dot plot generated is on side scatter (SSC-A, Y axis) and forward scatter (FSC-A, X axis). Gate on cell population in function of size-scatter allows to appreciate the singlet cells on the dot plot showing FSC-A (Y axis) and FSC-H (X axis). By gating on all singlet cell population, a dot plot representing SSC-A (Y axis) and viability marker (intensity mean, X axis) was generated, displaying cell viability by following the cells negative for the marker. Finally, by gating on all viable cell population, a dot plot representing Dapi (intensity mean, Y axis) and viral NP (intensity mean, X axis) was generated to quantify the infection per experimental condition. (C-E) Quantification of SARS-CoV-2 infection in full differentiated primary human lung organoid-derived ALI cultures per donor: (C) Histocytometry (left column, scatter plot with bar graph) based on Dapi and NP signal. (D) Flow cytometry (middle column, scatter plot with bar graph), based on cell viability and NP signal. (E) Viral titer (right column, scatter plot with bar graph) based on the number of plaques visualized by staining (see material and methods). Data shown are representative of at least 2 independent experiments per donor (four donors). Each replicate corresponds to viral titer, or the intensity mean percentage for histocytometry and flow cytometry data, represented by shape and color: Donor 1, white circle; donor 2 red square; donor 3 green triangle and donor 4 black diamond. Bars indicate mean.

**Figure S3 SARS-CoV-2 variants infection assessment of lung organoid-derived air-liquid interface (ALI) cultures by flow cytometry and plate assay (viral titers). (Related to figure 3)**

Quantification of SARS-CoV-2 infection in fully differentiated primary human lung organoid-derived ALI cultures per strain: (A) Flow cytometry (middle column, scatter plot with bar graph), based on cell viability and NP signal. (B) Viral titer (right column, scatter plot with bar graph) based on the number of plaques visualized by staining (see material and methods). Data shown are representatives of at least 2 independent experiments per strain. Each replicate corresponds to viral titer, or the intensity mean percentage for histocytometry and flow cytometry data, represented by shape and color: Donor 1, white circle; donor 2 red square; donor 3 green triangle and donor 4 black diamond. For strains: USA-WA1/2020: black, Beta: blue, Delta: red and Omicron: green. Bars indicate mean.

**Figure S4 Secretory and ciliated cells are predominantly infected by SARS-CoV-2 variants in primary human lung organoid-derived ALI cultures. (Related to figure 3)**

Representative immunofluorescent (IF) section (8 μm) of differentiated lung organoid-derived ALI cultures per variant (USA/WA1-2020 virus, Delta and Omicron tissue from donor 3 and Beta variant donor 4) at the peak of infection (reflected by NP expression), respectively, 6 days post infection (DPI) for USA/WA1-2020, 4 DPI for Beta and Delta, and 2 DPI for Omicron. Stained for epithelial cell subtypes: showing markers for basal cells (CK5, red), goblet cells (MUC5AC, red), club cells (SCGB1A1, white), ciliated cells (acetylated a-tubulin, white), Nuclei (Dapi, blue) and viral NP protein (green). Scale bar 20μm, in white on the left corner.

**Figure S5 GeoMX digital spatial profiling of SARS-CoV-2 (wuhan-like virus) infected ALI. (Related to figure 6)**

Sequencing quality is inspected for sufficient saturation, ensuring sensitivity of low expressors: (A) Plot of sequencing saturation percentage by ROI surface area ($um^2$). 78 ROIs were analyzed, 18,676 total targets. (B) Violin plots representing the data normalization to the third quartile (Q3) to account for differences in cellularity, ROI Size, etc. (C) Bar graph plot representing the 18,676 total target across the 78 ROIs, 0 samples below the 50% warning, 14,316 genes were expressed in 10% of ROIs and 1282 were expressed in 50 % of ROIs. (D) Heatmap representing the Q3 normalized counts expressed above LOQ in at least 95% of ROIs (17266 genes above LOQ) by slide-scan (time-points) and infection status. The sequencing was performed in two batches with at least 3 replicates per condition from two representative experiment with SARS-CoV-2 (donor 3 for slides #1, #2, #4 and #5; on slide #4, 2 sections from donor 4 and one from donor 3).

**Figure S6 CSF3 and CCL20 immunoregulators markers expression in ALI exposed to SARS-CoV-2 variants and in human lung epithelium with immune cells distribution. (Related to Figure 7)**

(A) Representative images of ALI (donor 3) mock-infected (controls media without virus at 6 days) and ALI infected with SARS-CoV-2 variants (105PFU) at 1, 3 and 6 days post infection (DPI) stained for nuclei (Dapi, blue), viral NP (white) to reveal the effective viral replication, CSF3 (green) and CCL20 (red). Scale bars 20 μm, in white on the left corner. (B) Representative zoomed images of human lung epithelium (Donor 1) stained for nuclei (Dapi, blue), CK5 (green), CSF3 (red), CCL20 (white) and Phalloidin (dark yellow). Scale bars 8 μm, in white on the left corner. (C) Representative images of human lung epithelium (Donor 1) stained for nuclei (Dapi, blue), CD11b (green), CD14 (magenta), CSF3 (white) and CCL20 (red). Scale bars 20 μm, in white on the left corner.

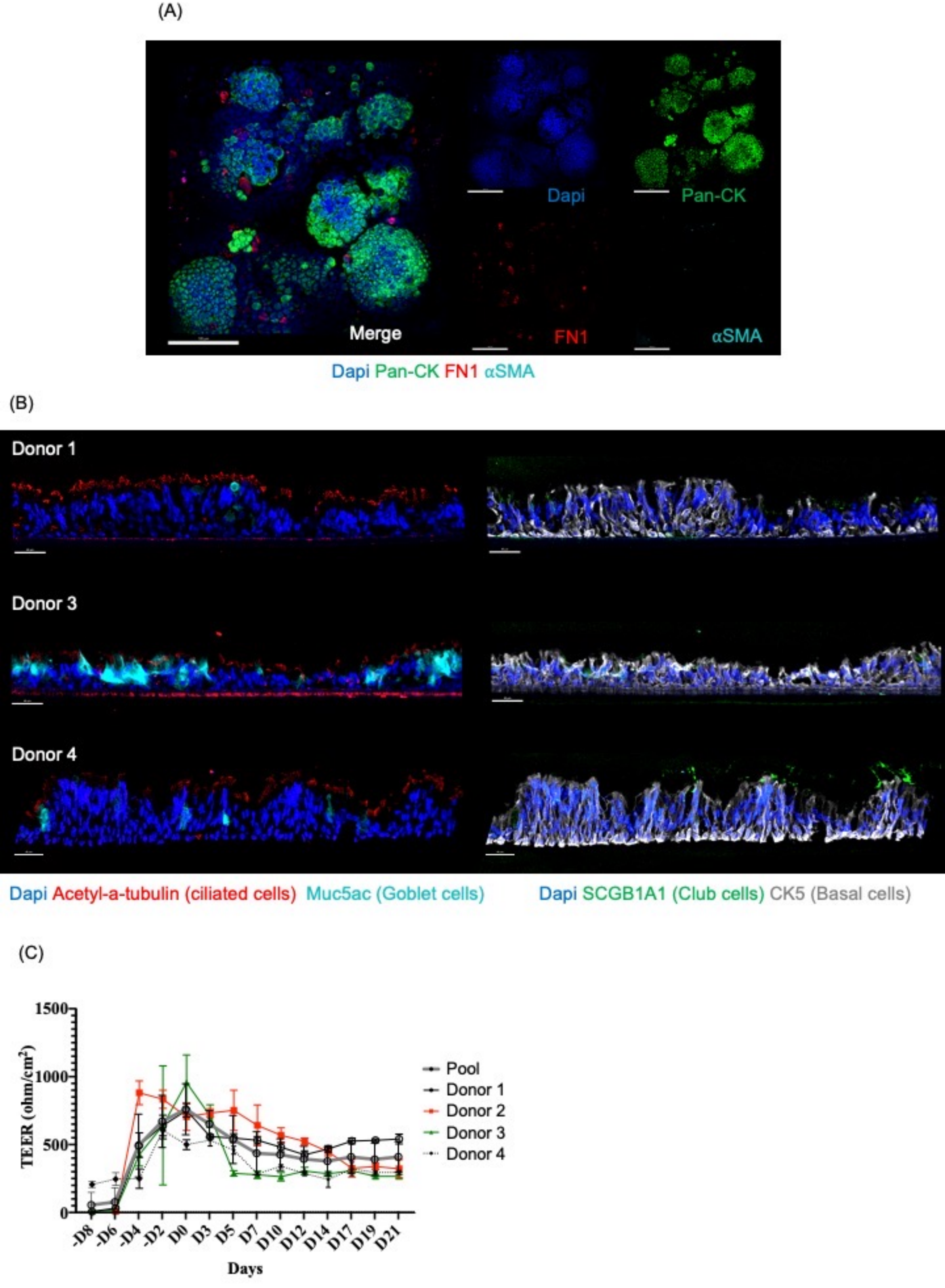
(A)
Dapi
Pan-CK
Merge
FN1
αSMA
Dapi Pan-CK FN1 αSMA
(B)
Donor 1
Donor 3
Donor 4
Dapi Acetyl-a-tubulin (ciliated cells) Muc5ac (Goblet cells)
Dapi SCGB1A1 (Club cells) CK5 (Basal cells)
(C)
TEER (ohm/cm²)
1500
1000
500
0
-D8
-D6
-D4
-D2
D0
D3
D5
D7
D10
D12
D14
D17
D19
D21
Days
Pool
Donor 1
Donor 2
Donor 3
Donor 4

**Figure S1**

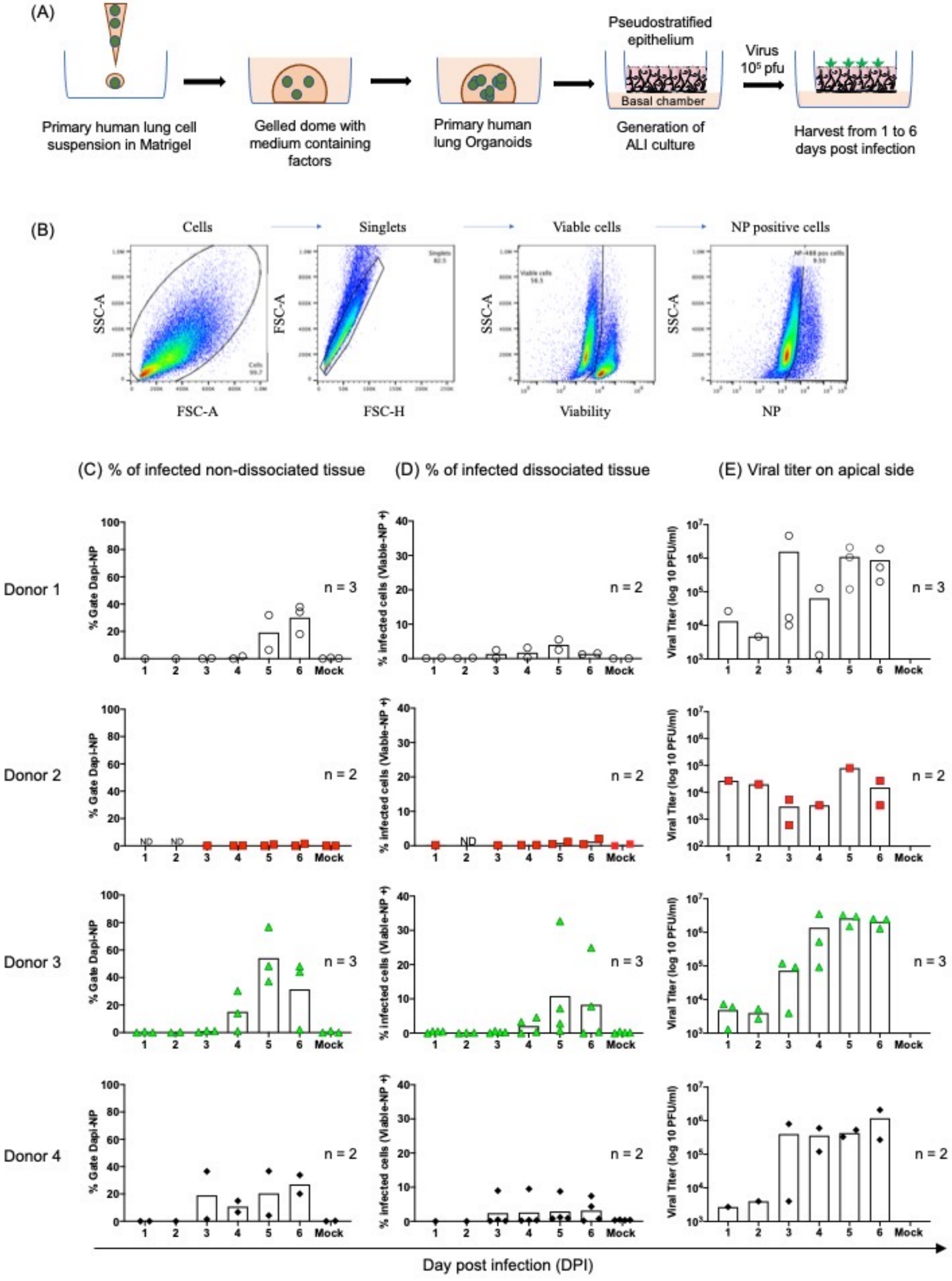

(A)
Primary human lung cell suspension in Matrigel
Gelled dome with medium containing factors
Primary human lung Organoids
Pseudostratified epithelium
Basal chamber
Generation of ALI culture
Virus
10⁵ pfu
Harvest from 1 to 6 days post infection
(B)
Cells
Singlets
Viable cells
NP positive cells
SSC-A
FSC-A
FSC-H
Viability
NP
(C) % of infected non-dissociated tissue
(D) % of infected dissociated tissue
(E) Viral titer on apical side
Donor 1
Donor 2
Donor 3
Donor 4
% Gate Dapi-NP
% infected cells (Viable-NP +)
Viral Titer (log 10 PFU/ml)
n = 3
n = 2
ND
Mock
Day post infection (DPI)


**Figure S2**

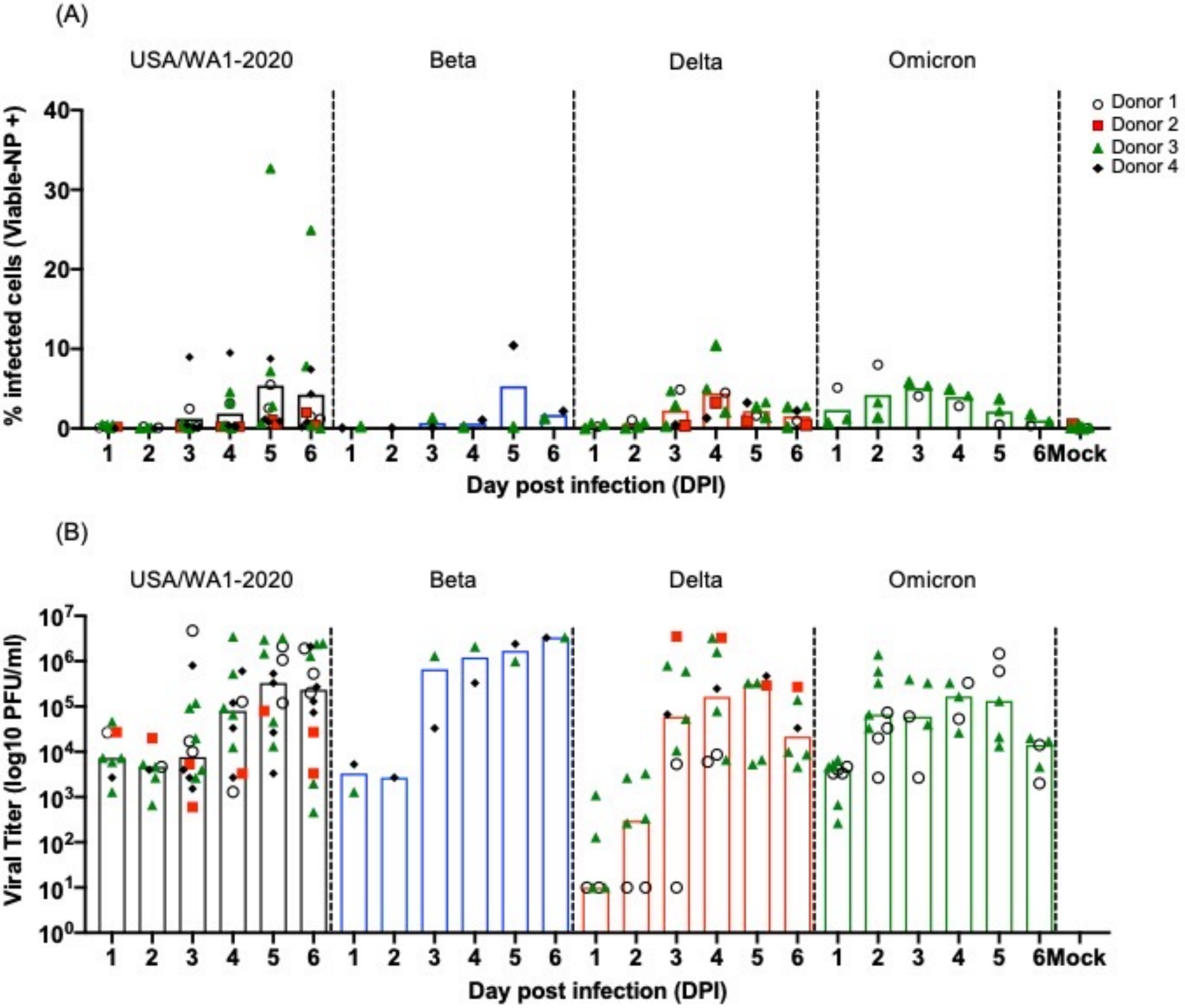
(A)
USA/WA1-2020
Beta
Delta
Omicron
% infected cells (Viable-NP +)
0
10
20
30
40
Donor 1
Donor 2
Donor 3
Donor 4
1
2
3
4
5
6
Mock
Day post infection (DPI)
(B)
Viral Titer (log10 PFU/ml)
$10^0$
$10^1$
$10^2$
$10^3$
$10^4$
$10^5$
$10^6$
$10^7$

**Figure S3**

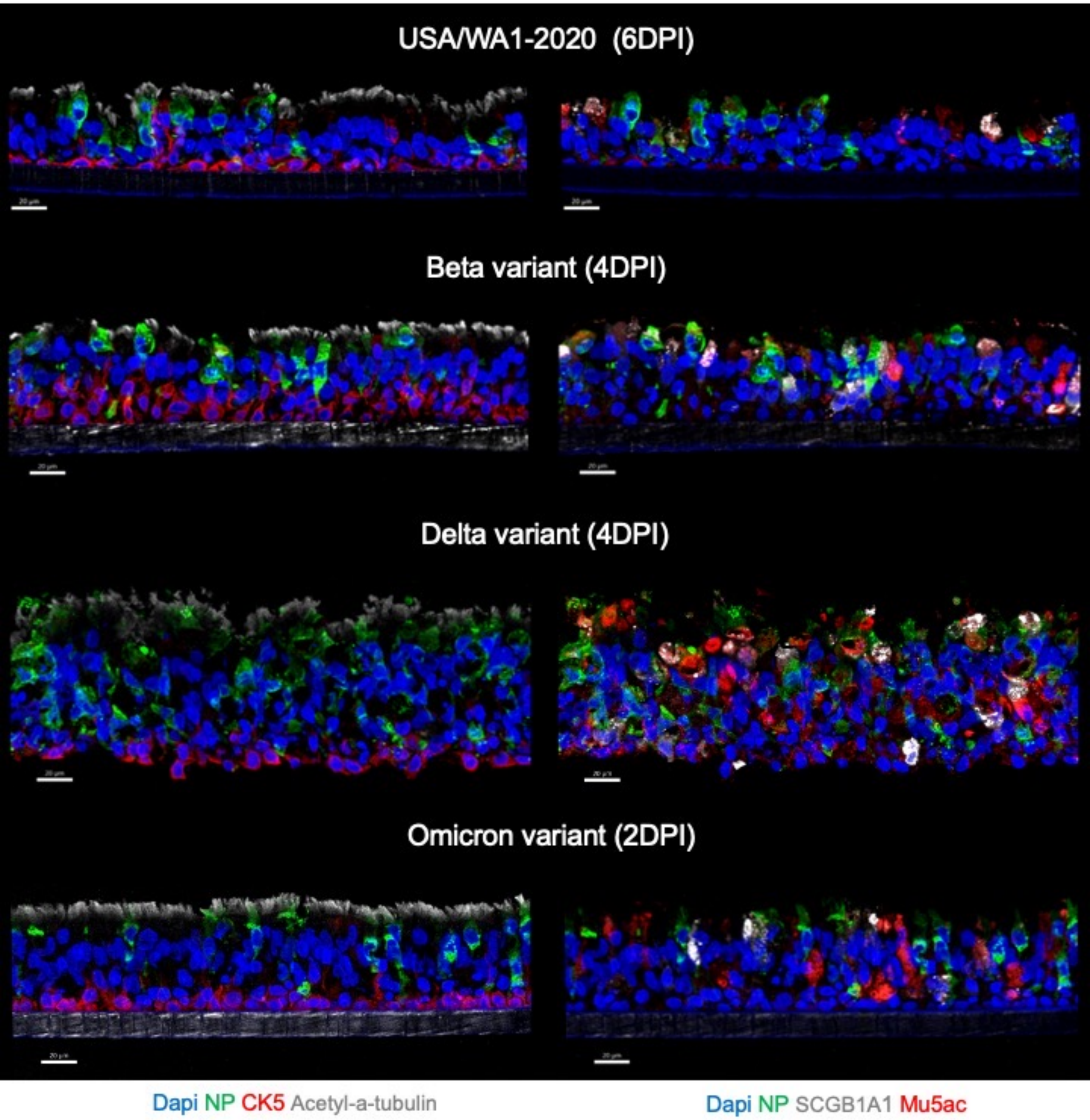

USA/WA1-2020 (6DPI)
Beta variant (4DPI)
Delta variant (4DPI)
Omicron variant (2DPI)
Dapi NP CK5 Acetyl-a-tubulin
Dapi NP SCGB1A1 Mu5ac


**Figure S4**

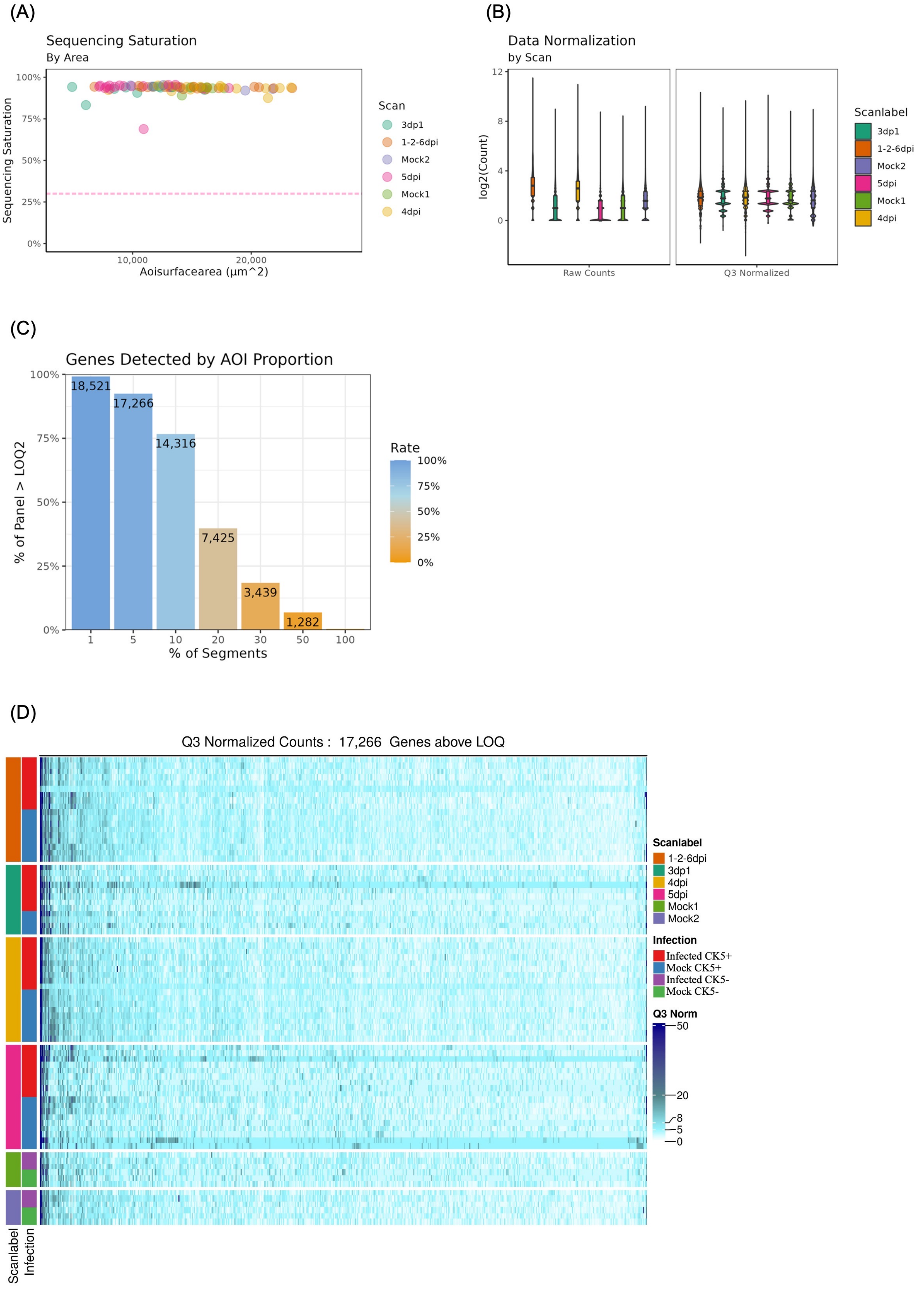

(A)
Sequencing Saturation
By Area
Sequencing Saturation
100%
75%
50%
25%
0%
10,000
20,000
Aoisurfacearea (µm^2)
Scan
3dp1
1-2-6dpi
Mock2
5dpi
Mock1
4dpi
(B)
Data Normalization
by Scan
log2(Count)
12
8
4
0
Raw Counts
Q3 Normalized
Scanlabel
3dp1
1-2-6dpi
Mock2
5dpi
Mock1
4dpi
(C)
Genes Detected by AOI Proportion
% of Panel > LOQ2
18,521
17,266
14,316
7,425
3,439
1,282
1
5
10
20
30
50
100
% of Segments
Rate
100%
75%
50%
25%
0%
(D)
Q3 Normalized Counts : 17,266 Genes above LOQ
Scanlabel
1-2-6dpi
3dp1
4dpi
5dpi
Mock1
Mock2
Infection
Infected CK5+
Mock CK5+
Infected CK5-
Mock CK5-
Q3 Norm
50
20
8
5
0
Scanlabel
Infection


**Figure S5**

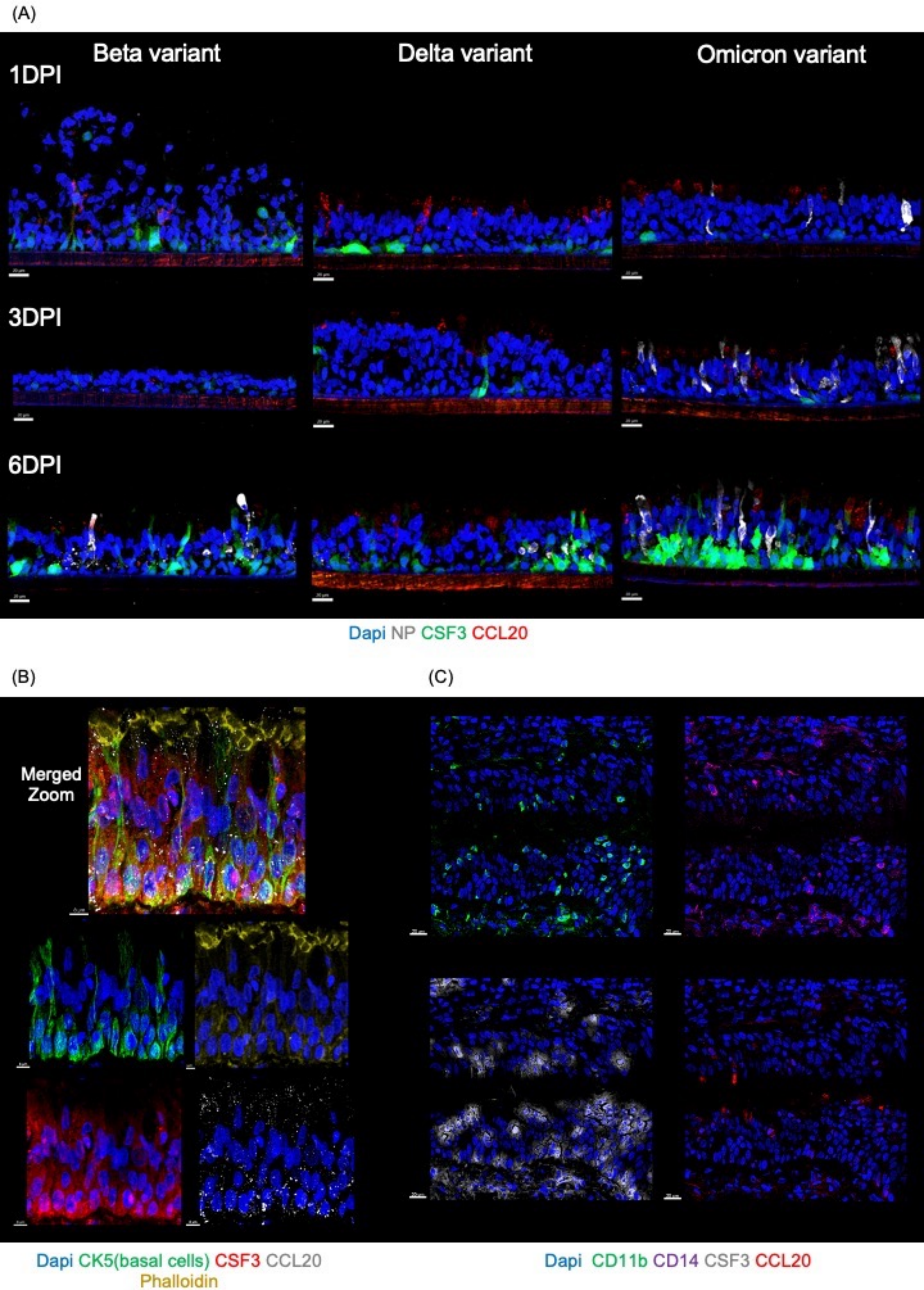
(A)
Beta variant
Delta variant
Omicron variant
1DPI
3DPI
6DPI
Dapi NP CSF3 CCL20
(B)
(C)
Merged Zoom
Dapi CK5(basal cells) CSF3 CCL20
Phalloidin
Dapi CD11b CD14 CSF3 CCL20

**Figure S6**
